# RegTools: Integrated analysis of genomic and transcriptomic data for the discovery of splice-associated variants in cancer

**DOI:** 10.1101/436634

**Authors:** Kelsy C. Cotto, Yang-Yang Feng, Avinash Ramu, Megan Richters, Sharon L. Freshour, Zachary L. Skidmore, Huiming Xia, Joshua F. McMichael, Jason Kunisaki, Katie M. Campbell, Timothy Hung-Po Chen, Emily B. Rozycki, Douglas Adkins, Siddhartha Devarakonda, Sumithra Sankararaman, Yiing Lin, William C. Chapman, Christopher A. Maher, Vivek Arora, Gavin P. Dunn, Ravindra Uppaluri, Ramaswamy Govindan, Obi L. Griffith, Malachi Griffith

## Abstract

Somatic mutations within non-coding regions and even exons may have unidentified regulatory consequences that are often overlooked in analysis workflows. Here we present RegTools (www.regtools.org), a computationally efficient, free, and open-source software package designed to integrate somatic variants from genomic data with splice junctions from bulk or single cell transcriptomic data to identify variants that may cause aberrant splicing. RegTools was applied to over 9,000 tumor samples with both tumor DNA and RNA sequence data. We discovered 235,778 events where a splice-associated variant significantly increased the splicing of a particular junction, across 158,200 unique variants and 131,212 unique junctions. To characterize these somatic variants and their associated splice isoforms, we annotated them with the Variant Effect Predictor (VEP), SpliceAI, and Genotype-Tissue Expression (GTEx) junction counts and compared our results to other tools that integrate genomic and transcriptomic data. While many events were corroborated by the aforementioned tools, the flexibility of RegTools also allowed us to identify novel splice-associated variants and previously unreported patterns of splicing disruption in known cancer drivers, such as *TP53, CDKN2A*, and *B2M*, as well as in genes not previously considered cancer-relevant.

## Introduction

Alternative splicing of messenger RNA allows a single gene to encode multiple gene products, increasing a cell’s functional diversity and regulatory precision. However, splicing malfunction can lead to imbalances in transcriptional output or even the presence of novel oncogenic transcripts^1^. The interpretation of variants in cancer is frequently focused on direct proteincoding alterations^2^. However, most somatic mutations arise in intronic and intergenic regions, and exonic mutations may also have unidentified regulatory consequences^3,4,5,6^. For example, mutations can affect splicing either in trans, by acting on splicing effectors, or in cis, by altering the splicing signals located on the affected pre-mRNA transcripts themselves^7^.

Increasingly, we are identifying the importance of cis-acting splice-associated variants in disease processes, including in cancer^8,9^. However, our understanding of the landscape of these variants is currently limited, and few tools exist for their discovery. One approach for identifying splice-associated variants has been to predict the strength of putative splice sites in pre-mRNA from genomic sequences, such as the method used by SpliceAI^10–13^. With the advent of efficient and affordable RNA-sequencing (RNA-seq), we are also seeing the development of tools that take the complementary approach of observing products of alternative splicing directly in RNA sequencing data, such as SUPPA2 and SPLADDER^14,15^. However, most of these tools have focused on the role of trans-acting splice-associated variants^16^. Only a few tools link products of alternative splicing to specific genomic variants to investigate their potential cis-acting role in splicing regulation, and these few tools have limitations that preclude them from broad applications. The sQTL-based approach taken by LeafCutter^17^ and others^18,19^ is designed for single-nucleotide polymorphisms, which occur with relatively high frequency, and is thus ill-suited to studying somatic variants, or any case in which the frequency of a particular variant is very low (often unique) in a given sample population. Recent tools created for large-scale analysis of cancer-specific data, such as MiSplice and Veridical, ignore certain types of alternative splicing, are tailored to specific analysis strategies and hypotheses, or are otherwise inaccessible to the end-user due to practical issues such as lack of documentation, difficulty with installation and integration with existing pipelines, limited computational efficiency, or license restrictions^20–22^. To address these needs, we have developed RegTools, a free, opensource (MIT license) software package that is well-documented, easy to use, and designed to efficiently and flexibly identify potential cis-acting splice-associated variants in tumors (www.regtools.org). At the highest level, RegTools contains three sub-modules: a *variants* module to annotate genomic variant calls for their potential splicing relevance, a *junctions* module to analyze aligned RNA-seq data to extract and annotate splice junctions, and a *cis-splice-effects* module that associates these variants and junctions to identify potential splice-associated variants. Each sub-module contains one or more commands, which can be used individually or integrated together to create customized splice-regulatory variant analysis pipelines.

To demonstrate the ability of RegTools to identify potential splice-associated variants from tumor data, we analyzed a combination of data available from the McDonnell Genome Institute (MGI) at Washington University School of Medicine and The Cancer Genome Atlas (TCGA) project. In total, we applied RegTools to 9,173 tumors across 35 cancer types. We compared RegTools with other tools that integrate genomic and transcriptomic data to identify potential splice-associated variants, specifically Veridical, MiSplice, and SAVNet^20,21,23^. Novel junctions identified by RegTools were compared to data from The Genotype-Tissue Expression (GTEx) project to assess whether these junctions are present in normal tissues^24^. Variants significantly associated with novel junctions were processed through VEP and Illumina’s SpliceAI tool to compare our findings against splicing consequences predicted based on variant information alone^13,25^. We identified variants in known cancer drivers, such as *TP53, CDKN2A*, and *B2M*, as well as in novel potential drivers, such as *RNF145*.

## Results

### The RegTools tool suite supports splice-associated variant discovery by the integration of genome and transcriptome data

RegTools is a tool suite composed of three modules designed to aid users in a broad range of splicing-related analyses. The *variants* module contains the *annotate* command. The *variants annotate* command takes a VCF of somatic variant calls and a GTF of transcript annotations as input. RegTools has no particular preference for variant callers or sources of reference transcript annotations. Each variant is annotated by RegTools with known overlapping genes and transcripts and is categorized into one of several user-configurable “variant types”, based on position relative to the edges of known exons. The variant type annotation depends on the stringency for splice-association that the user sets with the “splice variant window” setting. By default, RegTools marks intronic variants within 2 base pairs (bp) of the exon edge as “splicing intronic”, exonic variants within 3 bp as “splicing exonic”, other intronic variants as “intronic”, and other exonic variants as “exonic”. RegTools focuses on “splicing intronic” and “splicing exonic” in downstream analyses. To allow for the discovery of an arbitrarily expansive set of variants, RegTools allows the user to customize the size of the intronic/exonic windows individually (e.g. -i 2 -e 3 for default splice variant window, -i 50 -e 5 for intronic variants 50 bp from an exon edge and exonic variants 5 bp from an exon edge) or even consider all intronic/exonic variants as potentially splice-associated (e.g. -I or -E) (**Figure 1A**).

**Figure 1:**
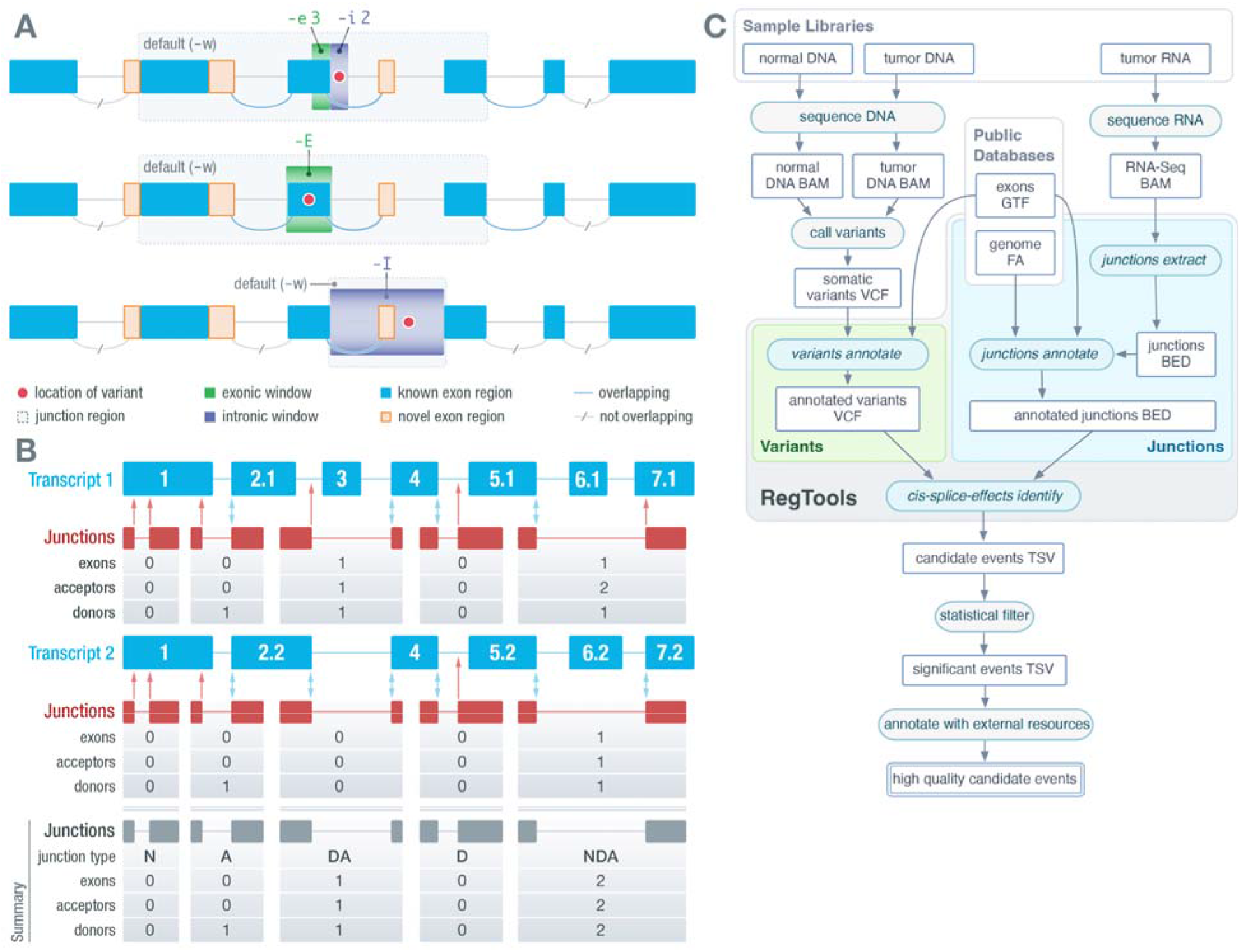
Regtools features individual modules and an integrated pipeline for flexible, streamlined discovery of cis-acting splice-associated variants. **A)** By default, *variants annotate* marks variants within 3 bp on the exonic side and 2 bp on the intronic side of an exon edge as potentially splice-associated. This “splice variant window” can be modified individually for the exonic side and intronic side using the “-e” and “-i” options, respectively. With *cis-splice-effects identify*, for each variant in the splice variant window, a “splice junction region” is determined by finding the largest span of sequence space between exons that flank the exon associated with the splicing-relevant variant. The splice junction region can also be set manually to contain the entire sequence space of *n* bases upstream and downstream of the variant using the “-w” option. Junctions overlapping the splice junction region are associated with the variant. Using the -E option considers all exonic variants as potentially splice-associated, but is otherwise the same. Using the -I option considers all intronic variants as potentially splice-associated and also limits the splice junction region to the intronic region in which the variant is found, excluding the flanking exons. **B)** *Cis-splice-effects identify* and the underlying *junctions annotate* command annotate junctions based on whether the donor and acceptor site combination is found in the reference transcriptome GTF. In this example, there are two known transcripts (shown in blue) that overlap a set of junctions observed in RNAseq data (depicted as junction supporting reads in red). Comparing the observed junctions to the reference junctions in the first transcript (top panel), RegTools checks to see if the observed donor and acceptor splice sites are found in any of the reference exons and also counts the number of exons, acceptors, and donors skipped by a particular junction. Double blue arrows represent matches between observed and reference donor/acceptor sites, while single red arrows show novel splice sites. These steps are repeated for the rest of the relevant transcripts, keeping track of whether there are known donor/acceptor combinations. Junctions with a known donor but novel acceptor or vice-versa are annotated as “D” or “A”, respectively. If both sites are known but do not appear in combination in any transcripts, the junction is annotated as “NDA”, whereas if both sites are unknown, the junction is annotated as “N”. If the junction is known to the reference GTF, it is marked as “DA”. **C)** The *cis-splice-effects identify* command relies on the *variants annotate*, *junctions extract*, and *junctions annotate* submodules. This pipeline takes variant calls and RNA-seq alignments along with genome and transcriptome references and outputs information about events (pairs of variants and associated junctions). RegTools is agnostic to downstream research goals, and its output can be filtered through user-specific methods and thus can be applied to a broad set of scientific questions.

The *junctions* module contains the *extract* and *annotate* commands. The *junctions extract* command takes a BAM/CRAM file containing aligned RNA-seq reads, infers the exon-exon boundaries based on the CIGAR strings^26^, and outputs each “junction” as a feature in BED12 format. The *junctions annotate* command takes a BED file containing junctions in BED12 format (such as the one produced by *junctions extract*), a FASTA file containing the reference genome, and a GTF file containing reference transcriptome annotations and generates a TSV file, annotating each junction with: the number of acceptor sites, donor sites, and exons skipped, and the identities of known overlapping transcripts and genes. We also annotate the “junction type”, which denotes if and how the junction is novel (i.e. not found in the reference transcriptome). If the donor site is known, but the acceptor site is not or vice-versa, it is marked as “D” or “A”, respectively. If both the donor and acceptor sites are known, but the pairing is not known, it is marked as “NDA”. If both the donor and acceptor sites are unknown, it is marked as “N”. If the junction is not novel (i.e. it appears in at least one transcript in the supplied GTF), it is marked as “DA” (**Figure 1B**).

The *cis-splice-effects* module contains the *identify* and *associate* commands, which identify potential splice-associated variants from genomic and transcriptomic data. The *cis-splice-effects identify* command requires the following files as input: a VCF file containing variant calls, an alignment file containing aligned RNA-seq reads, a reference genome FASTA file, and a reference transcriptome GTF file. The *identify* pipeline internally relies on *variants annotate, junctions extract*, and *junctions annotate* to output a TSV containing junctions proximal to putatively splice-associated variants. The *identify* pipeline can be customized using the same parameters as in the individual commands. Briefly, *cis-splice-effects identify* first performs *variants annotate* to determine the splicing relevance of each variant in the input VCF. For each variant, a “splice junction region” is determined by finding the largest span of sequence space between exons that flank the variant-containing exon. From here, *junctions extract* identifies splicing junctions present in the RNA-seq alignment. Next, *junctions annotate* labels each extracted junction with information from the reference transcriptome as described above and its associated variants based on splice junction region overlap (**Figure 1C**). To enable the association of variants with pre-extracted junctions, *cis-splice-effects associate* performs the same pipeline as *cis-splice-effects identify*, but takes junctions from an existing BED12 file, such as one previously created by the *junction extract* command, instead of re-extracting from the alignment file.

For our analysis, we annotated the pairs of variants and associated junctions identified by RegTools, which we refer to as “events”, with additional information such as whether this association was identified by a comparable tool, whether the junction was found in GTEx, and whether the event occurred in a cancer gene according to the Cancer Gene Census (CGC) (**Figure 1C**)^24,27^. Finally, for each event identified by RegTools, we created an IGV session that showed a BED file with the junction, a VCF file with the variant, and BAM files with DNA alignments for all samples that contained the variant^28^. These IGV sessions were used to manually review candidate events to assess whether the association between the variant and junction was biologically plausible.

Overall, RegTools is designed for broad applicability and computational efficiency. By relying on well-established and widely adopted standards for sequence alignments (BAM/CRAM), annotation files (GTF), and variant calls (VCF) and by remaining agnostic to downstream statistical methods and comparisons, our tool can be applied to a broad set of scientific queries and datasets. Moreover, performance tests show that *cis-splice-effects identify* can process a typical candidate variant list of 1,500,000 variants and a corresponding RNA-seq BAM file of 82,807,868 reads in just ~8 minutes (**Supplementary Figure 1**). Run time increases approximately linearly with increasing numbers of junctions and variants.

### Pan-cancer analysis of 35 tumor types identifies somatic variants that alter canonical splicing

RegTools was applied to 9,173 samples over 35 cancer types. 32 of these cohorts came from TCGA while the remaining 3 were obtained from other projects being conducted at MGI. Cohort sizes ranged from 21 to 1,022 samples. In total, 6,370,631 somatic variants (**Supplementary Figure 2A**) and 2,387,989,201 junction observations (**Supplementary Figure 2B**) were analyzed by RegTools. By comparing the number of initial variants to the number of statistically significant variants, we see that RegTools produces a highly prioritized list of potential splice-associated variants **(Supplementary Figure 3)**. Additionally, when analyzing the junctions within each sample, we found that known junctions present in the reference transcriptome are frequently seen within GTEx data while novel junctions are rarely seen within GTEx **(Supplementary Figure 4)**. These represent potential tumor-specific junctions. We identified 235,778 significant events for novel junctions that use a known donor and novel acceptor (D), novel donor and known acceptor (A), or novel combination of a known donor and a known acceptor (NDA) (**Methods**, **Supplementary Figure 2C, Supplementary Files 1 and 2**). Additionally, we identified 5,157 events for known (DA) junctions (**Supplementary Files 3 and 4**). Thus, while splice-associated variants usually result in a novel junction occurring, they may also alter the relative amounts of known junctions. Generally, significant events were evenly distributed among the novel junction types considered (D, A, and NDA). The number of significant events increased as the splice variant window size increased, with both the E and I results being comparable in number. Notably, hepatocellular carcinoma (HCC) was the only cohort that had whole genome sequencing (WGS) data available and, as expected, it exhibited a marked increase in the number of significant events for its results within the “I” splice variant window. This observation highlights the low sequence coverage of intronic regions that occurs with whole exome sequencing (WES), which reduces the potential for the discovery of splice-associated variants within introns.

Variants were analyzed across tumor types for how often each resulted in either single or multiple novel junctions (**Figure 2A**). While variants were most commonly associated with a single novel junction (72.3-83.8%), they could also be associated with multiple junctions, either of the same type (6.6-10.9%) or of different types (9.7-16.8%) (**Figure 2B**). Variants that are associated with multiple novel junctions of different types were further investigated to identify how often a particular junction type occurred with another (**Figure 2C**). Most commonly, variants were associated with either novel donor or acceptor site usage (A or D) and with an exon-skipping junction (NDA). These kinds of events were particularly common within the default window (2 intronic bases or 3 exonic bases from the exon edge), potentially due to variants within these positions having a high probability of disrupting the natural splice site, thus causing the splicing machinery to use a cryptic splice site nearby or skip the exon entirely. The next most common co-occurrence was a variant being associated with both novel donor site usage leading to A junctions and acceptor site usage leading to D junctions. The occurrence of a variant associated with the combination of a novel donor, novel acceptor, and exon-skipping was low, and remained low, even as the search space increased with the larger splice variant windows. Overall, this analysis highlights that there is evidence that a single variant can lead to multiple novel junctions being expressed. Tools such as SpliceAI only allow for a single junction to be associated with a variant and therefore may not completely describe the splicing effects of the variant in question for up to ~27% of cases.

**Figure 2.**
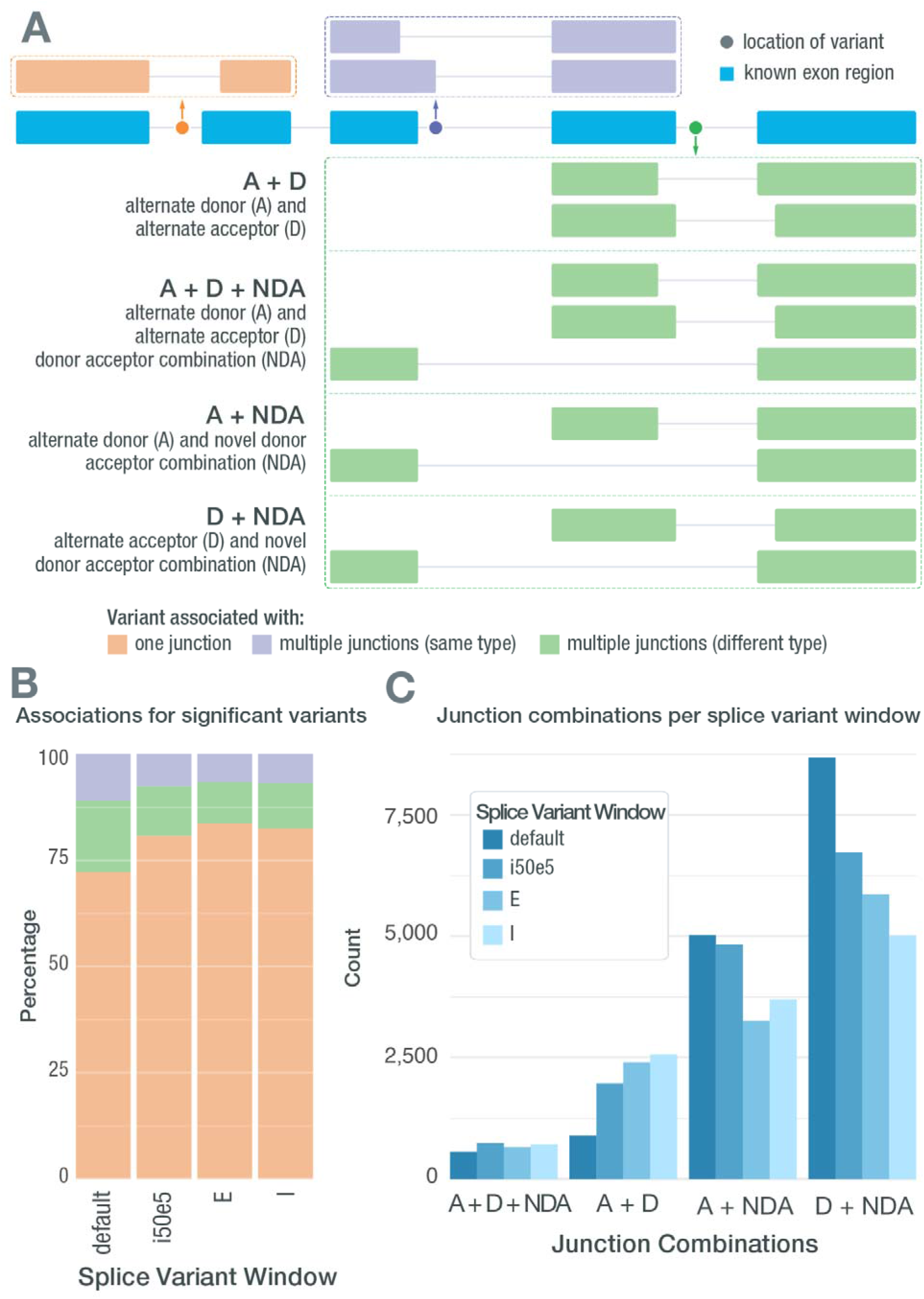
Splice-associated variants may result in multiple novel junctions. **A)** A single splice-associated variant can result in a single novel junction, multiple novel junctions of the same junction type, or multiple novel junctions of different junction types. Depicted is a variant resulting in a single novel junction (orange), a variant resulting in two novel junctions that both use alternate donor sites (purple), and a variant resulting in multiple junctions of different types (green). **B)** Stacked bars showing how often significant splice-associated variants are associated with only one junction, multiple junctions of the same type, or multiple junctions of different types. **C)** Bar chart showing how often each junction combination occurs when a single splice-associated variant results in multiple junctions of different types in each of the RegTools splice variant windows used.

### Orthogonal validation of RegTools using clinical data and verified splice-associated variants

We tested RegTools against multiple datasets to further validate this tool suite. The first dataset that we compared against was the 10 splice-site-creating variants that Jayasinghe et al. validated using mini-gene functional assays^20^. They selected 11 variants that their tool, MiSplice, originally identified from TCGA data. These mutations were then compared to wildtype sequences using a pCAS2.1 splicing reporter mini-gene functional assay and 10 were validated through sequencing of alternatively spliced products. These 10 variants were run through RegTools using corresponding aligned transcriptomic reads for each sample. RegTools identified an association between all 10 variants and an aberrant splice junction **(Supplementary File 5)**.

The next dataset that we used to validate RegTools was MutSpliceDB^29^. This is a public database that contains manually reviewed RNA-based evidence of the effects of splice site variants on splicing. Currently, data is curated from TCGA and the Cancer Cell Line Encyclopedia^30–32^. When we accessed MutSpliceDB, there were 211 entries. Out of these 211 entries, 208 were annotated by MutSpliceDB as either intron inclusion or exon skipping events. We used the mutations provided and the corresponding RNA alignments to process each of these mutations through RegTools. We detected all 211 manually reviewed splice site variants **(Supplementary File 6)**.

We also validated RegTools using clinical sequencing projects that allowed us to directly test the effects of somatic variants between multiple tumors within individuals. The first dataset utilized is from Schaettler and Richters et al. (2022) which investigated the impact of spatial heterogeneity on genomic characteristics of gliomas and brain metastases^33^. For this study, tumor tissue was surgically resected from 30 patients. Immediately following resection, each sample was dissociated into multiple (2-4) spatially distinct tumor regions that then underwent WES and RNA sequencing. We ran RegTools to identify splice-associated variants within each distinct tumor region. A benefit of the heterogeneity of these samples and the multisector approach that was used is that we were able to interrogate many examples of clonal and subclonal splice variants. This allowed us to validate associations within other tumor regions based on whether the variant was also present within those regions. Through this approach, we validated 134 out of 146 splice-associated variants in samples where multiple sectors shared the same variant and aberrant junction. Conversely, we found 142 splice-associated variants out of 212 in which one sector contained a variant and novel splice junction but other regions in which both the variant and associated junction were absent **(Supplementary File 7)**. In other words, the events predicted by RegTools in the RNA data reflected the spatial heterogeneity of somatic mutations observed in the DNA. This provides a form of biological validation that is more representative of true splicing biology than a typical mini-gene assay approach.

Another dataset that we employed for a conceptually similar biological validation was treatment-matched naive and post-treatment recurrence samples of small cell lung cancer (SCLC)^34^. By applying RegTools to these samples, we found splice-associated variants that persisted from the treatment-naive sample to the recurrence sample (0%-36.0%). Additionally, we identified samples where a splice-associated variant was lost due to treatment or arose post-treatment, either through the growth of a previously existing subclone or the emergence of a novel splice-associated variant (64.0-100%) **(Supplementary File 8)**. In this analysis, the RegTools results reflected the temporal heterogeneity of the tumors under treatment.

We also validated RegTools using long-read sequencing data to confirm the full-length structure of alternatively spliced isoforms inferred from short read data. For this analysis, we used a well-described breast cancer cell line, HCC1395. For a normal comparator, we used HCC1395’s matched lymphoblastoid cell line, HCC1395BL. For each of these samples, whole genome, exome and RNA-seq were performed. For HCC1395, Oxford Nanopore Technologies long-read sequencing was performed using both the Direct RNA Sequencing Kit and Direct cDNA Sequencing Kit. After applying RegTools to the bulk genomic and transcriptomic data and obtaining candidate splice-associated variants, we validated 80% of novel junctions observed within the short-read data and confirmed the resulting novel transcript sequences **(Supplementary File 9)**.

Finally, we validated RegTools on a single-cell RNA (scRNA) sequencing dataset from a study investigating the mechanisms of response to immune checkpoint blockade (ICB) using MCB6C, a transplantable organoid model of urothelial carcinoma with features of human basal-squamous urothelial carcinoma^35^. This model had also been subjected to WES of DNA isolated from tumor cells and matched normal cells from the tail of the mouse originally used to create the tumor. Analysis of the tumor/normal WES DNA was performed to identify somatic variants. We then identified single cells from three conditions and surveyed their expressed transcripts for evidence of the somatic variants. Each cell was then classified as either tumor or normal, based on somatic variant expression, and separated into corresponding alignment files. More specifically, to identify a tumor cell, we used the following criteria: two or more somatic variants detected with >20X total coverage, >5 variant reads, and >10% variant allele fraction (VAF). To identify a normal cell, we used the following criteria: no variants detected and two or more of the variant positions with >20X total coverage. Using these criteria, we defined 5,587 tumor cells and 17,022 normal cells for a total of 22,609 single cells. We processed these cells through an updated version of RegTools modified to support single-cell data, treating each cell as an individual sample. This approach allowed us to greatly increase our power for determining tumor-associated splice-associated variants due to all mutations being tumor-specific and each cell representing an independent readout of the splicing machinery. We were able to identify over 300 splice-associated variants that had multiple cells of support, including within *Trp53* and *Bin1* **(Figure 3, Supplementary Figure 5)**. Within *Trp53*, we identify an intronic variant (mm10, chr11:g.69589711T>G; c.1067+2 position of intron 8 of transcript NM_011640.3) that is associated with the skipping of exon 8. This exon contains important domains such as binding domains for DNA and *Axin* in addition to a bipartite nuclear localization signal (UniProt: P02340)^36,37^. Similarly, we identify an intronic variant in *Bin1* (mm10, chr18:g.32432427T>C, c.1516+2 position of intron 14 of transcript NM_001083334.1) that is associated with an alternate donor site being used and partial retention of intron 14. *Bin1* has been shown to have tumor suppressor properties and shown to be dysregulated in breast cancer, neuroblastoma, prostate cancer, and melanoma^38–42^. The ability to identify such events at a single cell resolution may provide insights into how splice-associated variants contribute to tumorigenesis and tumor progression in ways that are not possible through bulk sequencing approaches.

**Figure 3:**
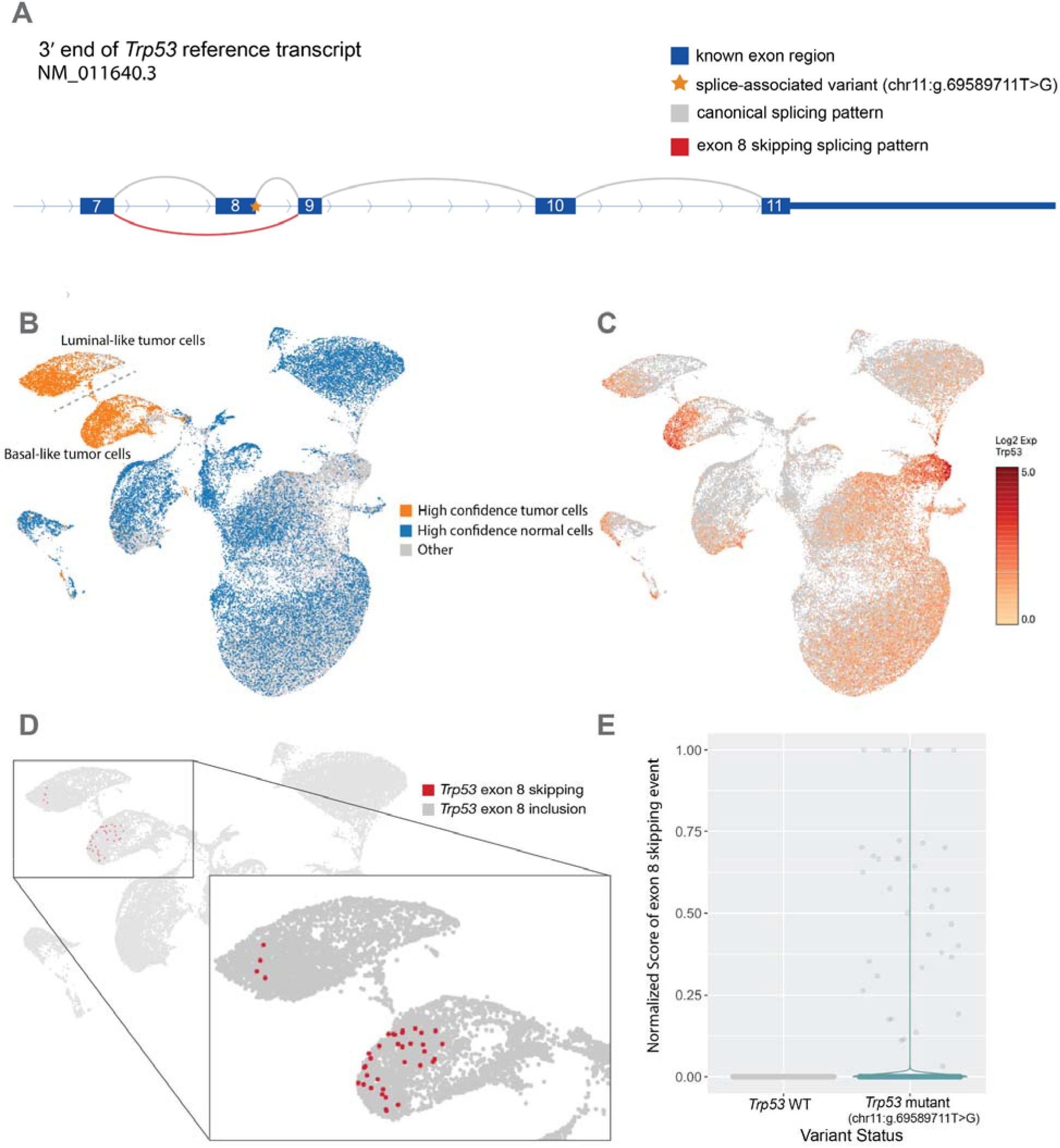
Intronic SNV in *Trp53* associated with exon 8 skipping. **A)** Schematic of a single nucleotide splice donor variant (mm10, chr11:g.69589711T>G; c.1067+2 position of intron 8 of transcript NM_011640.3) within intron 8 of *Trp53*. This variant is significantly associated (p < 0.001) with an exon skipping event causing the formation of an NDA junction. This result was found using the default splice variant window parameter (i2e3). **B)** UMAP projection of single cells from MCB6C organoid derived tumors with high confidence tumor cells (orange) and high confidence normal cells (blue) highlighted. **C)** UMAP projection of single cells from MCB6C organoid derived tumors overlaid with Log2 expression of Trp53. **D)** Zoomed view of cells containing the Trp53 exon skipping event. **E)** Violin plot comparing the normalized junction score of the novel exon skipping event in cells with and without the variant.

Through the application of RegTools to the aforementioned datasets, we were able to identify high-quality, validated splice-associated variants. Additionally, we utilized well-designed clinical and scRNA datasets to identify tumor-specific splice-associated mutations more stringently. These results demonstrate the broad utility of RegTools and its ability to identify splice-associated somatic variants robustly.

### Pan-cancer analysis reveals novel splicing patterns within known cancer genes and potential cancer drivers

While efforts have been made to associate variants with specific cancer types, there has been little focus on identifying cancer-specific splicing variants, even those in known cancer genes. *TP53* is a rare example of a driver whose splice-associated variants are well-characterized in numerous cancer types^43^. To investigate the impact of variants on splicing disruption in cancer genes across different cancer types, we further analyzed significant events to identify genes that had recurrent splice-associated variants. Within each cohort, we looked for recurrent genes using two separate metrics: a binomial test p-value and the fraction of samples (see Methods). For ranking and selecting the most recurrent genes, each metric was computed by pooling across all cohorts. For assessing cancer-type specificity, each metric was then also computed using only results from a given cancer cohort. Since the mechanisms underlying the creation of novel junctions versus the disruption of existing splicing patterns may be different, analysis was performed separately for D/A/NDA junctions **(Figure 4, Supplementary Files 10-13)** and DA junctions **(Supplementary Figure 6, Supplementary File 14)**, which allowed multiple test correction in accordance with the noise of the respective data. We identified 6,954 genes in which there was at least one variant predicted to influence the splicing of a D/A/NDA junction. The 99th percentile of these genes, when ranked by either metric, is significantly enriched for known cancer genes, as annotated by the CGC (p=1.26E-19, ranked by binomial p-values, p=2.97E-24, ranked by the fraction of samples; hypergeometric test). We also identified 3,643 genes in which there was at least one variant predicted to influence the splicing of a DA (known) junction. The 99th percentile of these genes, when ranked by either metric, is also significantly enriched for known cancer genes, as annotated by the CGC (p=1.00E-04, ranked by binomial p-values, p=3.56E-07, ranked by the fraction of samples; hypergeometric test). We also performed the same analyses using either the TCGA or MGI cohorts alone. The TCGA-only analyses gave similar results to the combined analyses, with the 99th percentile of genes found in the D/A/NDA and DA analyses again enriched for cancer genes **(Supplementary Figures 7 and 8)**. Due to small cohort sizes, in the MGI-only analyses, we identified only 329 and 208 genes in the D/A/NDA and DA analyses, respectively. The 99th percentile of genes was not significantly enriched for cancer genes in either of these analyses **(Supplementary Figures 9 and 10)**.

**Figure 4.**
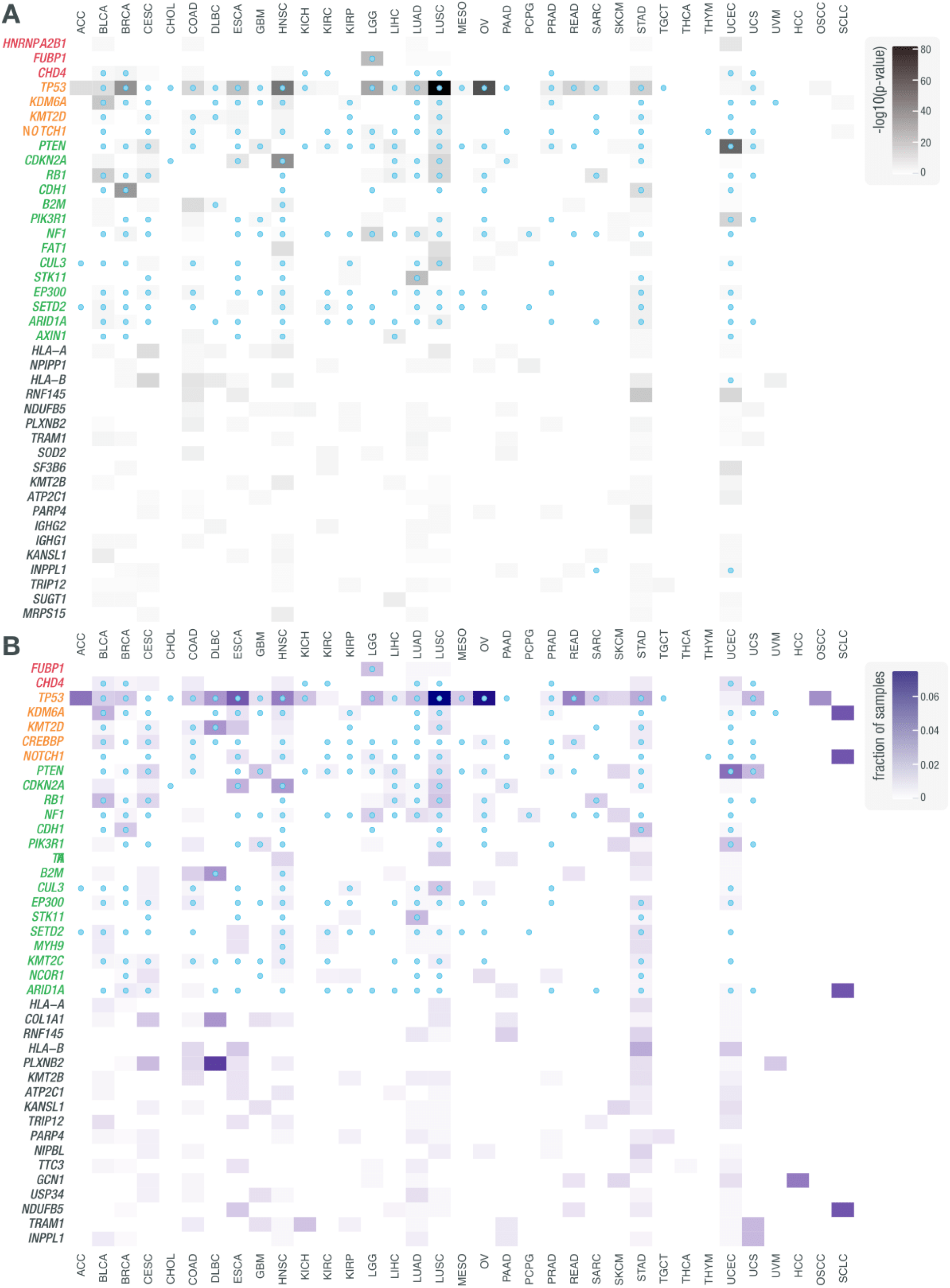
Pan-cancer analysis of cohorts from TCGA and MGI reveals genes recurrently disrupted by variants that are associated with non-canonical splicing patterns. Results of analysis for recurrently disrupted genes in each cohort. **A)** Rows correspond to the 40 most frequently recurring genes, as ranked by binomial p-value across cohorts. Genes are clustered by whether they were annotated by the CGC as an oncogene (red), an oncogene and tumor suppressor gene (yellow), a tumor suppressor gene (green), or another type of cancer-relevant gene. Shading corresponds to -log10(p value) and columns represent cancer types. Red marks within cells indicate that the gene was annotated by CHASMplus as a driver within a given TCGA cohort. **B**) Rows correspond to the 40 most frequently recurring genes, as ranked by the fraction of samples across cohorts. Shading corresponds to the fraction of samples and columns represent cancer types. Blue dots within cells indicate that the gene was annotated by CHASMplus as a driver within a given TCGA cohort. These results were obtained using the default splice variant window parameter (i2e3).

When analyzing D, A, and NDA junctions, we saw an enrichment for known tumor suppressor genes among the most splice-disrupted genes, including several examples where splice disruption is a known mechanism such as *TP53, PTEN, CDKN2A*, and *RB1*. Specifically, in the case of *TP53*, we identified 428 variants that were significantly associated with at least one novel junction. One such example is the intronic SNV (GRCh38, chr17:g.7673609C>A) that was identified in an OSCC sample and was associated with exon skipping and novel acceptor site usage, with 23 and 41 reads of support, respectively **(Supplementary Figure 11)**. The cancer types in which we find splice disruption of *TP53* and other known cancer genes is in concordance with associations between genes and cancer types described by CGC and CHASMplus^27,44^. Our identification of known drivers, many with known susceptibilities to splicing dysregulation in cancer, indicates the ability of our method to identify true splicing effects that are likely cancer-relevant. Additional splice-associated variants were found in genes not currently known to be linked to cancer. Some of these genes, such as *IGHG1* and *IGHG2*, are located in regions of the genome with high genetic variability in the population and are at loci where the reference genome may not represent structural diversity. These regions tend to result in false positive somatic variant calls and misalignment of short reads. These factors will complicate the identification of true splice regulatory variants in these regions. These regions also undergo V-D-J recombination in B cells, and some aligned reads could correspond to DNA from infiltrating immune cells. Some studies exclude immune-related regions of the genome entirely because of these kinds of complexities^20^. However, disruption of these genes may still be relevant to tumor biology and certainly tumor immunotherapy ^45–48^.

Another cancer gene that had a recurrence of splice-associated variants was *B2M*. Specifically, we identified six samples with intronic variants on either side of exon 2 **(Figure 5)**. These mutations were identified by VEP to be either splice acceptor or splice donor variants and were also identified by Veridical. SpliceAI identified one of the novel junctions for each variant but failed to identify additional novel junctions, as SpliceAI only identifies one novel acceptor and donor site per variant. Notably, 4 out of the 6 samples that these variants were found in are Microsatellite instability-high (MSI-H) tumors^49^. Mutations in *B2M*, particularly within colorectal MSI-H tumors, have been identified as a method for tumors to disable HLA class I antigen-mediated presentation^50^. Furthermore, in a study of patients treated with immune checkpoint blockade (ICB) therapy, defects affecting B2M were observed in 29.4% of patients with progressing disease^51^. In the same study, *B2M* mutations were exclusively seen in pretreatment samples from patients who did not respond to ICB or in post-progression samples after the initial response to ICB^51^. There are several genes responsible for the processing, loading, and presentation of antigens that are mutated in cancers^52^. However, no proteins can be substituted for B2M in HLA class I presentation, thus making the loss of B2M a particularly robust method for ICB resistance^53^. We also observed exonic variants and variants further in intronic regions that may disrupt canonical splicing of B2M. These findings raise the possibility that intronic variants may enable tumor immune escape by disrupting B2M splicing.

**Figure 5.**
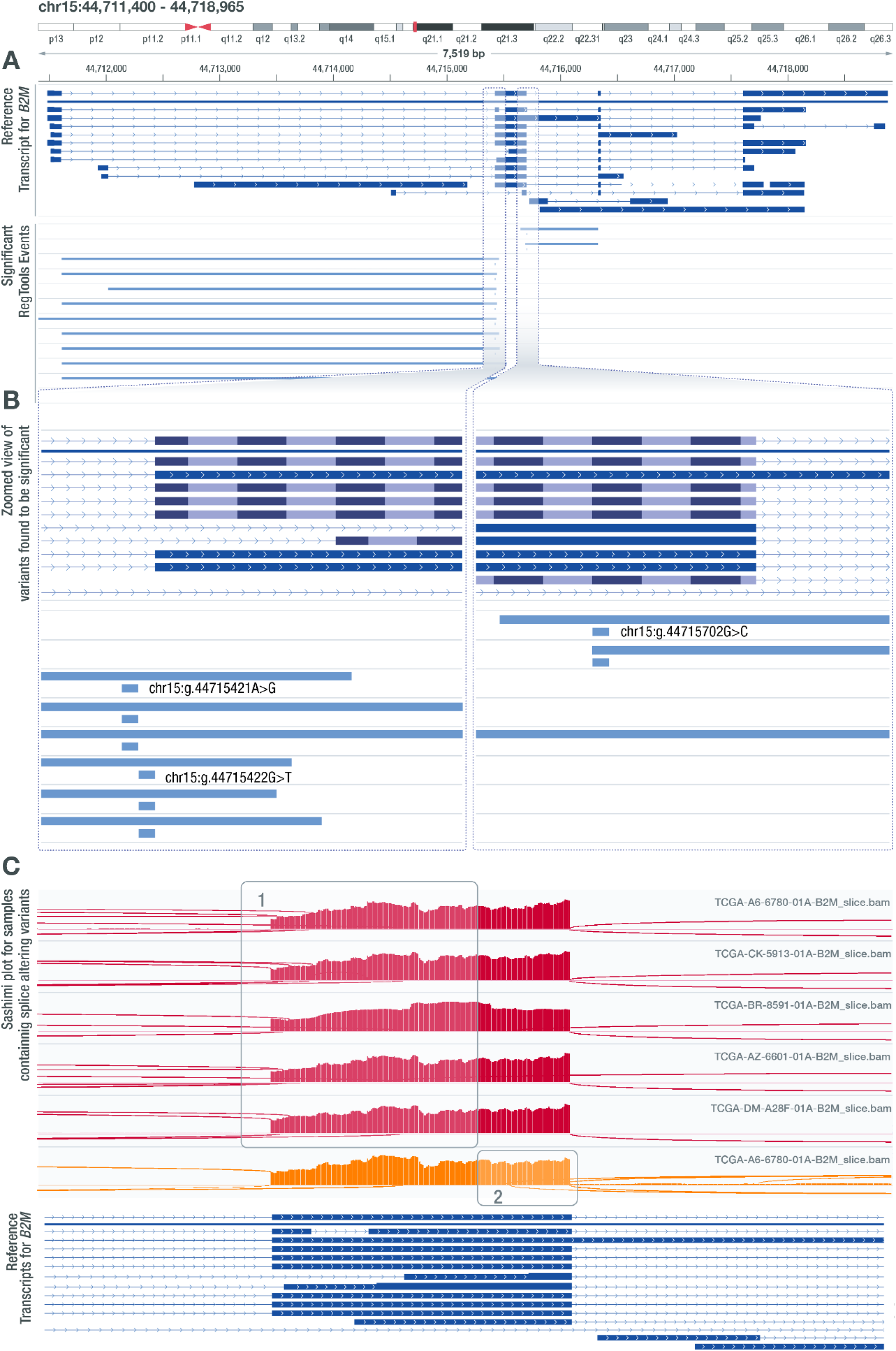
Several SNVs in *B2M* are associated with alternate acceptor and alternate donor usage. **A)** IGV snapshot of three intronic variant positions (GRCh38 - chr15:g.44715421A>G, chr15:g.44715422G>T, chr15:g.44715702G>C) found to be associated with alternate acceptor and donor usage that leads to the formation of novel transcript products. This result was found using the default splice variant window parameter (i2e3). **B)** Zoomed in view of the variants identified by RegTools that are associated with alternate acceptor and donor usage. Two of these variant positions flank the acceptor site and one variant flanks the donor site of the area that is being affected. **C)** Sashimi plot visualizations for samples containing the identified variants that show 1) alternate acceptor usage (red) or 2) alternate donor usage (orange).

We also identified recurrent splice-associated variants in genes not currently known to be cancer genes (according to CGC), such as *RNF145*. RegTools identified a recurrent single base pair deletion that results in the skipping of exon 8 (**Supplementary Figure 12**). This gene is a paralog of *RNF139*, which is mutated in several MSI-H cancer types^54^. This event was found in STAD, UCEC, COAD, and ESCA tumors, all of which are considered to be MSI-H tumors^49^. Analyzing the effect of the skipping of exon 8 on the mRNA sequence, we observed that the reading frame remains intact, possibly leading to a gain of function event. Additionally, the skipping of exon 8 leads to the removal of a transmembrane domain and a phosphorylation site, S352, which could be important for the regulation of this gene^55^. Based on these findings, splicing disruption of *RNF145* warrants further investigation as a potential driver mechanism underlying MSI-H cancers.

While most of our analysis focused on splice-associated variants that resulted in novel junctions, we also investigated variants that shifted the relative amounts of known junctions. We identified several variants that led to alternate donor usage in *CDKN2A*, a key tumor suppressor gene^56^ (**Supplementary Figure 13)**. When these variants are present, an alternate known donor site is used that leads to the formation of the transcript ENST00000579122.1 instead of ENST00000304494.9, the transcript that encodes for p16^ink4a^, a known tumor suppressor. The transcript that results from the use of this alternative donor site is missing the last twenty-eight amino acids that form the C-terminal end of p16^ink4a^. Notably, this removes two phosphorylation sites within the p16 protein, S140 and S152, which could disrupt the association of p16^ink4a^ with CDK4^57^. This highlights the importance of including known transcripts in alternative splicing analyses, as variants may alter splice site usage in a way that results in a known, but still potentially oncogenic transcript product.

### RegTools provides usability and flexibility in integrating genomic and transcriptomic data to identify splice-associated variants

To evaluate the performance of RegTools, we compared our results to those of SAVNet, MiSplice, Veridical, VEP, and SpliceAI^13,20,21,23,25^. These tools vary in their inputs and methodology for identifying splice-associated variants (**Figure 6A**). Like RegTools, SAVNet, MiSplice, and Veridical integrate genomic and transcriptomic data to identify splice-associated variants and have also been utilized in pan-cancer analyses that have demonstrated the utility of this integrative approach. However, there are practical and methodological limitations of these tools that impede their broad application. MiSplice and Veridical have varying levels of code availability or portability. MiSplice is available via GitHub as a collection of Perl scripts built to run via Load Sharing Facility (LSF) job scheduling. To run MiSplice without an LSF cluster, code changes are required. Veridical is only available via a subscription through CytoGnomix’s MutationForecaster. Similar to RegTools, SAVNet is available via GitHub or a Docker image. However, unlike Regtools, SAVNet relies on splicing junction files generated by STAR^58^ whereas RegTools can use RNA-seq alignment files from HISAT2^59^, TopHat2^60^, or STAR, thus allowing it to be easily integrated into bioinformatics workflows that use any of these popular aligners or to use pre-existing alignments. To demonstrate the time needed to generate the STAR splicing junction files in order to run SAVNet, we benchmarked RegTools and SAVNet using LUAD samples from TCGA **(Supplementary Figure 14)**. On average, Regtools was 3.2x faster, taking into account the unalignment and realignment SAVnet required to generate the necessary starting files from STAR. Moreover, these tools prescribe certain analytical and methodological frameworks, whereas Regtools is designed to offer greater usability and flexibility to control how genomic and transcriptomic data is integrated. SAVNet, MiSplice, and Veridical employ particular statistical methods for the identification of splice-associated variants, whereas Regtools can be integrated at any step in the pipeline. Additionally, some of these tools filter out any transcripts found within the reference transcriptome, precluding the investigation of canonical splicing patterns as can be done by examining DA junctions with RegTools, and do not allow the user to set a custom window in which they wish to focus splice-associated variant discovery (e.g. around the splice site, all exonic variants, etc.). Furthermore, MiSplice does not include exon-skipping events. RegTools addresses these limitations by identifying what pieces of information to extract from a sample’s genome and transcriptome in a basic, easily configurable way that allows for generalization.

**Figure 6.**
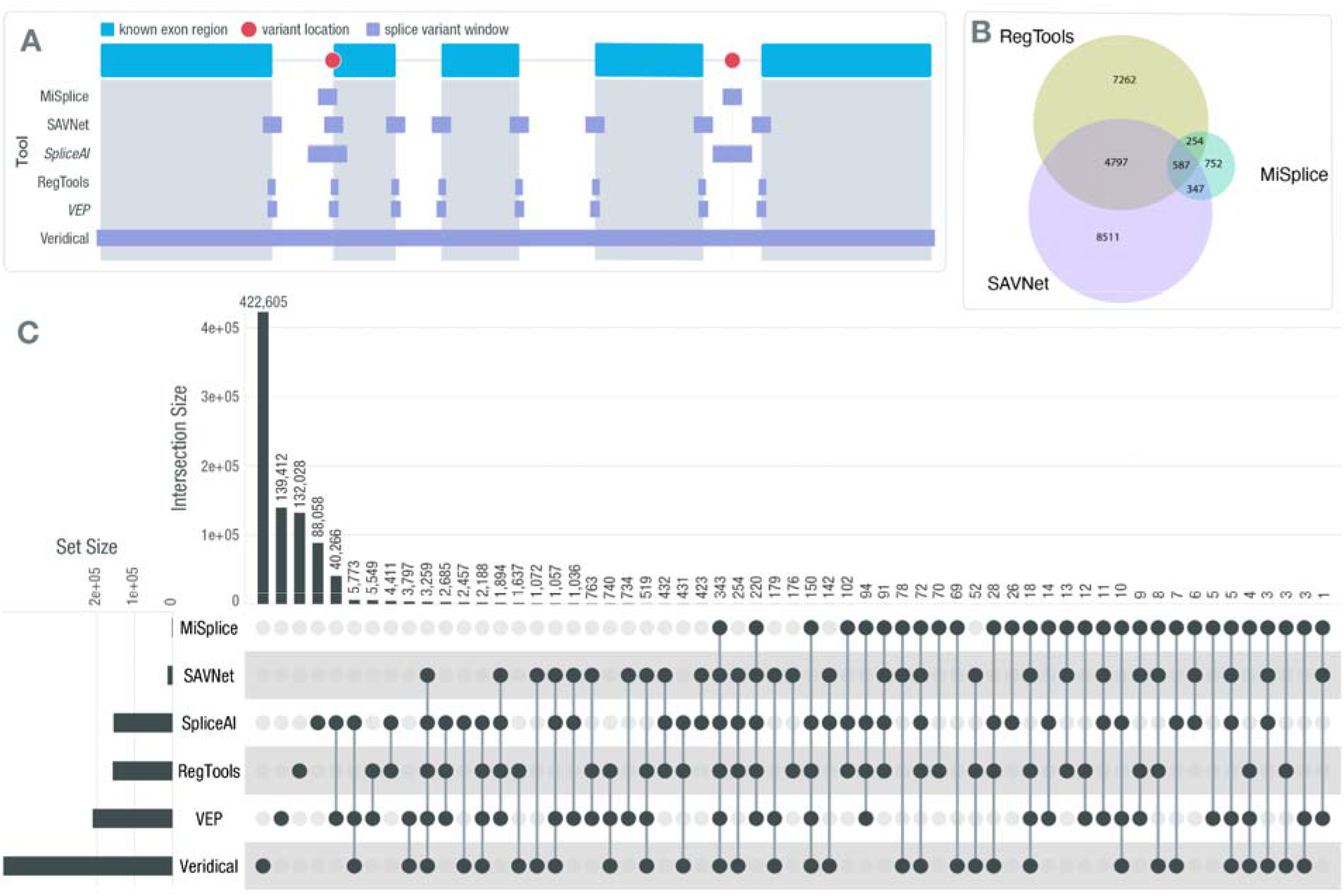
Comparison of RegTools with other tools for identifying cis-acting splice-associated variants. **A)** Conceptual diagram of contrasting approaches employed by various tools for identifying cis-acting splice-associated variants. For this example, the splice variant window for RegTools is its default splice variant window employed for our main analyses. An *italicized* tool name indicates that the tool only considers genomic data for making its calls, instead of a combination of genomic and transcriptomic data. **B)** Venn diagram comparing the splice-associated variants identified by RegTools, using its default splice window parameter, MiSplice, and SAVNet. **C)** UpSet plot comparing splice-associated variants identified by RegTools both the -E and -I splice variant window parameters to those identified by other splice variant predictors and annotators using their default settings. Each tool’s total number of variant predictions is shown on the left sidebar graph. The number of variants specific to each tool or shared between different combinations of tools is indicated by the bar graph along the top, with the individual or connected dots indicating the tools.

The set of splice-associated variants identified using Regtools with its default splice variant window (-i 2 -e 3) are most similar to MiSplice and SAVNet. These three result sets contain fewer splice-associated variants compared to Veridical due to the more tightly constrained search space for variants to be associated with splicing alterations. Thus, we primarily focused our comparison to MiSplice and SAVNet (**Figure 6B**). Compared to Regtools and SAVNet, MiSplice finds fewer splice-associated variants, which could be due to MiSplice not examining exon skipping junctions, starting with only the subset of variants in the Multi-Center Mutation Calling in Multiple Cancers (MC3) MAF^61^, and limiting junctions to those within 20 bp of the variant. However, MiSplice also detected many splice-associated variants that were not detected by Regtools or SAVNet, which could be due to these tools focusing on variants only within a limited distance from exon edges (**Figure 6A**). The concordance between Regtools and SAVNet was relatively high, and their respective concordances with MiSplice were comparable. These results illustrate that distinct methodologies will lead to distinct findings, which will be necessary to address the manifold goals and challenges of studying cis-splicing regulation. Focusing on variants closer to the exon edge may lead to a higher rate of true discoveries, given the established mechanistic relationship between splice site disruption and alternative splicing. However, there are also more distal splice regulatory elements, such as splicing enhancers and silencers or genomic sequences that resemble splice site motifs that can have effects on splicing patterns. Therefore, one may wish to expand the genomic area in which to identify splice-associated variants. An example that illustrates the importance of this is the identification of several breast cancer samples that have splice-associated variants within *GATA3* by RegTools. In our i50e5 analysis, we detected a set of highly recurrent *GATA3* mutations. Specifically, when focusing on mutations that occur within the RegTools splice variant window of i50e5 but outside the default window, we found 20 samples that contained the same 2 bp deletion (rs763236375), with 19 of these samples having alternative donor site usage for exon 5 of *GATA3* leading to a frameshifted protein product that lacks one of two zinc finger DNA binding domains. Within these samples, the novel junction becomes the dominant splice product compared to the canonical splice junction. Interestingly, this is a highly tumorspecific event, with this splice-associated variant only being found within breast cancer (**Supplementary File 11**). *GATA3* is a transcription factor, and its expression in breast cancer strongly correlates with estrogen receptor (ER) expression. This gene is mutated in approximately 10-15% of breast cancer cases, suggesting these are driver mutations, and during progression to metastatic breast cancer, GATA3 expression decreases^62,63^. These results highlight the value of a tool such as Regtools, which offers methodological flexibility to meet the diverse goals and challenges of studying splicing regulation.

In their recent publications, SAVNet^23^, MiSplice^20^, and Veridical^21,22^ also analyzed data from TCGA, with only minor differences in the number of samples included for each study. We also compared the results of these studies with the results obtained by RegTools when expanding the set of variants to include all exonic and intronic space. In this comparison, Veridical and RegTools identify a large number of splice-associated variants (**Figure 6C**). While this approach is the least biased, it is undoubtedly hindered by specificity and multiple testing challenges. This is due not just to the larger number of candidates, but also to the biology of splicing regulation - the density of true cis-regulatory splicing elements is not uniform in the genome and is, for example, higher around exon edges^64^. While we do know that some splicing regulatory elements such as exonic splicing silencers (ESSs), exonic splicing enhancers (ESEs), intronic splicing silencers (ISSs), and intronic splicing enhancers (ISEs) can be quite distal^65–67^, running RegTools or any tool in a mode that is capable of detecting these certainly creates a signal to noise challenge and will lead to candidate event lists with a higher false positive rate. Still, the identification of these distal splicing regulatory sequences and variants that modify their effects will ultimately be required to fully uncover the underlying mechanisms of diseases, such as cancer.

Both VEP and SpliceAI only consider information about the variant and its genomic sequence context and do not consider information from a sample’s transcriptome. A variant is considered to be splice-associated according to VEP if it occurs within 1-3 bases on the exonic side or 1-8 bases on the intronic side of a splice site. SpliceAI does not have restrictions on where the variant can occur in relation to the splice site, but by default, it predicts one new donor and acceptor site within 50 bp of the variant, based on reference transcript sequences from GENCODE. VEP and SpliceAI results were obtained by running each tool on all starting variants for the 35 cohorts included in this study. SpliceAI and VEP called a large number of variants either alone or in agreement with each other that none of the tools that integrate transcriptomic data from samples identified (**Figure 6C**). This indicates the limited reliability of approaches that make predictions based on genomic data alone without interrogating sample matched transcriptomic data, particularly in a disease context featuring novel junctions.

## Discussion

Splice-associated variants are often overlooked in traditional genomic analysis. Of the tools that exist, some only analyze genomic data, focus on junctions where either the canonical donor or acceptor site is affected (missing junctions that result from complete exon skipping), or consider only those variants within a narrow distance from known splice sites. To address these limitations, we created RegTools, a software suite for the analysis of variants and junctions in a splicing context. By relying on well-established standards for analyzing genomic and transcriptomic data and allowing flexible analysis parameters, we enable users to apply RegTools to a wide set of scientific methodologies and datasets. RegTools can include any kind of junction type, including exon-exon junctions that have ends that are not known donor/acceptor sites according to the GTF file (N junction according to RegTools), and any splice variant window size. To facilitate the use, portability and integration of RegTools into analysis workflows, we provide documentation and example workflows via (regtools.org) and provide a Docker image with all necessary software installed.

In order to demonstrate the utility of our tool, we applied RegTools to 9,173 tumor samples across 35 tumor types to profile the landscape of this category of variants. From this analysis, we report 133,987 variants that are associated with novel junctions that were missed by VEP or SpliceAI. We found splice-associated variants beyond the splice site consensus sequence, shift transcript usage between known transcripts, or result in novel exon-exon junctions. Specifically, we describe notable findings within *B2M*, *CDKN2A, and RNF145*. These results demonstrate the utility of RegTools in discovering putative splice-associated variants and confirm the importance of integrating RNA and DNA sequencing data in understanding the consequences of somatic mutations in cancer. To allow for validation and further investigation of these identified events, we make all of our annotated result files (**Supplementary Files 1-4**) and recurrence analysis files (**Supplementary Files 10-14**) available.

For certain RegTools results, such as those from E and I splice variant windows, there are higher numbers of splice-associated variants identified because of the broader region of consideration. One must be careful in comparing these results to other tools that have a more focused region of consideration. The increased number of events identified by RegTools in these comparisons does not necessarily suggest poor sensitivity of the other tools, but rather reflects that RegTools is being run in a mode that casts a wider net in order to identify more distal splice-associated variants, such as those in distal splice regulatory elements. This consideration highlights and reinforces that RegTools is highly configurable, and certain parameters that one can modify will impact sensitivity and specificity. For users that are concerned with avoiding false positives and less worried about maximizing sensitivity, we provide guidance on best practices for use of RegTools via our documentation at regtools.org. This includes the type of alignments RegTools supports, how to set the region of consideration, which junction types to focus on (e.g., NDA, DA, etc.), how to interpret the statistics results, recommended count thresholds, how to annotate with supporting information from GTEX, SpliceAI, and VEP, and much more. Because of the versatility and modularity of RegTools, we believe that it can be implemented into a variety of bioinformatics workflows to aid in the processing of sequencing data in disease studies or to answer specific questions about splicing biology.

Understanding the splicing landscape is crucial for unlocking potential therapeutic avenues in precision medicine and elucidating the basic mechanisms of splicing and cancer progression. The exploration of novel tumor-specific junctions will undoubtedly lead to translational applications, from discovering novel tumor drivers, diagnostic and prognostic biomarkers, and drug targets, to identifying a previously untapped source of neoantigens for personalized immunotherapy. While our analysis focused on splice-associated variants in cancer, we believe RegTools will play an important role in answering a broad range of questions across different disease states and biological processes by helping users extract splicing information from transcriptome data and linking it to somatic or germline variant calls. The computational efficiency of RegTools and the increasing availability of genomic and transcriptomic datasets will enable the investigation of splice regulatory motifs that have proven difficult to define such as exonic and intronic splicing enhancers and silencers. Any group with paired DNA and RNA-seq data stands to benefit from the functionality of RegTools.

## Methods

### Software implementation

RegTools is written in C++. CMake is used to build the executable from the source code. We have designed the RegTools package to be self-contained to minimize external software dependencies. A Unix platform with a C++ compiler and CMake is the minimum prerequisite for installing RegTools. Documentation for RegTools is maintained as text files within the source repository to minimize divergence from the code. We have implemented common file-handling tasks in RegTools with the help of open-source code from Samtools/HTSlib^26^ and BEDTools^68^ in an effort to ensure fast performance, consistent file handling, and interoperability with any aligner that adheres to the BAM specification. Statistical tests are conducted within RegTools using the RMath framework. GitHub actions and Coveralls are used to automate and monitor software compilation and unit tests to ensure software functionality. We utilized the Google Test framework to write unit tests.

RegTools consists of a core set of modules for variant annotation, junction extraction, junction annotation, and GTF utilities. Higher-level modules such as *cis-splice-effects* use the lower level modules to perform more complex analyses. We hope that bioinformaticians familiar with C/C++ can re-use or adapt the RegTools code (released under the open source MIT license) to implement similar tasks.

### Benchmarking

Performance metrics were calculated for all RegTools commands. Each command was run with default parameters on a single blade server (Intel(R) Xeon(R) CPU E5-2660 v2 @ 2.20GHz) with 10 GB of RAM and 10 replicates for each data point **(Supplementary Figure 1)**. Specifically for *cis-splice-effects identify*, we started with random selections of somatic variants, ranging from 10,000-1,500,000, across 8 data subsets. Using the output from *cis-splice-effects identify, variants annotate* was run on somatic variants from the 8 subsets (range: 0-17,742) predicted to have a splicing consequence. The function *junctions extract* was performed on the HCC1395 tumor RNA-seq data aligned with HISAT to GRCh37 and randomly downsampled at intervals ranging from 10-100%. Using output from *junctions extract, junctions annotate* was performed for 7 data subsets ranging from 1,000-500,000 randomly selected junctions.

Benchmark tests revealed an approximately linear performance for all functions. Variance between real and CPU time is highly dependent on the I/O speed of the write-disk and could account for artificially inflated real-time values given multiple jobs writing to the same disk simultaneously. The most computationally expensive function in a typical analysis workflow was *junctions extract*, which on average processed 33,091 reads/second (CPU) and took an average of 43.4 real vs 41.7 CPU minutes to run on a full bam file (82,807,868 reads total). The function *junctions annotate* was the next most computationally intensive function and took an average of 33.0 real/8.55 CPU minutes to run on 500,000 junctions, processing 975 junctions/second (CPU). The other functions were comparatively faster with *cis-splice-effects identify* and *variants annotate* able to process 3,105 and 118 variants per second (CPU), respectively. To process a typical candidate variant list of 1,500,000 variants and a corresponding RNA-seq BAM file of 82,807,868 reads with *cis-splice-effects identify* takes ~ 8.20 real/8.05 CPU minutes **(Supplementary Figure 1)**.

Performance metrics were also calculated for the statistics script and its associated wrapper script that handles dividing the variants into smaller chunks for processing to limit RAM usage. This command, *compare_junctions*, was benchmarked in January 2020 using Amazon Web Services (AWS) on a m5.4xlarge instance, based on the Amazon Linux 2 AMI, with 64 Gb of RAM, 16 vCPUs, and a mounted 1 TB SSD EBS volume with 3,000 IOPS. These data were generated from running *compare_junctions* on each of the included cohorts, with the largest being our BRCA cohort (1,022 samples) which processed 3.64 events per second (CPU).

For the benchmarking comparison between RegTools and SAVNet, we utilized fifty LUAD samples from TCGA. For our comparison, we imagined a use case where an individual would start by downloading alignment files from the Genomic Data Commons (GDC) Data Portal. For RegTools CPU and real-time measurements, *regtools junctions extract, regtools cis-splice-effects associate*, and *compare_junctions* were run for each sample. For SAVNet’s CPU and real-time measurements, alignment files were first unaligned using *SamToFastq* and then realigned using STAR to get each sample’s splice junction file, which is unavailable from the GDC Data Portal. Following these steps, SAVNet was then run and the time was added to that from the unalignment and realignment step. On average, it took SAVNet 3.2 times (real-time) as long as RegTools to run on the same samples when considering the unalignment and realignment required to generate the necessary starting files **(Supplementary Figure 14)**.

### Using RegTools to identify cis-acting, splice-associated variants

RegTools contains three sub-modules: “variants”, “junctions”, and “cis-splice-effects”. For complete instructions on usage, including a detailed workflow for how to analyze cohorts using RegTools, please visit regtools.org.

#### Variants annotate

This command takes a list of variants in VCF format. The file should be gzipped and indexed with Tabix^69^. The user must also supply a GTF file that specifies the reference transcriptome used to annotate the variants.

The INFO column of each line in the VCF is populated with comma-separated lists of the variant-overlapping genes, variant-overlapping transcripts, the distance between the variant and the associated exon edge for each transcript (i.e. each start or end of an exon whose splice variant window included the variant) defined as *min*(distance_from_start_of_exon, distance_from_end_of_exon), and the variant type for each transcript.

Internally, this function relies on HTSlib to parse the VCF file and search for features in the GTF file which overlap the variant. The splice variant window size (i.e. the maximum distance from the edge of an exon used to consider a variant as splice-associated) can be set by the options “-e <number of bases>” and “-i <number of bases>” for exonic and intronic variants, respectively. The variant type for each variant thus depends on the options used to set the splice variant window size. Variants captured by the window set by “-e” or “-i” are annotated as “splicing_exonic” and “splicing_intronic”, respectively. Alternatively, the “-E” and “-I” options can be used to analyze all exonic or intronic variants. These options do not change the variant type annotation, and variants found in these windows are labeled simply as “exonic” or “intronic”. By default, single exon transcripts are ignored, but they can be included with the “-S” option. By default, output is written to STDOUT in VCF format. To write to a file, use the option “-o <PATH/TO/FILE>”.

#### Junctions extract

This command takes an alignment file containing aligned RNA-seq reads and infers junctions (i.e. exon-exon boundaries) based on skipped regions in alignments as determined by the CIGAR string operator codes. These junctions are written to STDOUT in BED12 format. Alternatively, the output can be redirected to a file with the “-o <PATH/TO/FILE>”. RegTools ascertains strand information based on the XS tags set by the aligner, but can also determine the inferred strand of transcription based on the BAM flags if a stranded library strategy was employed. In the latter case, the strand specificity of the library can be provided using “-s <INT>” where 0 = unstranded, 1 = first-strand/RF, 2 = second-strand/FR. We have tested RegTools with alignment files from HISAT2^59^, TopHat2^60^, STAR^58^, kallisto^70^, or minimap2^71^, though we recommend HISAT2 or STAR for short read data and minimap2 for long read data. We have tested RegTools with data from the following sequencing platforms: Illumina, Oxford Nanopore Technologies, and 10X Genomics.

Users can set thresholds for minimum anchor length and minimum/maximum intron length. The minimum anchor length determines how many contiguous, matched base pairs on either side of the junction are required to include it in the final output. The required overlap can be observed amongst separated reads, whose union determines the thickStart and thickEnd of the BED feature. By default, a junction must have 8 bp anchors on each side to be counted but this can be set using the option “-a <minimum anchor length>”. The intron length is simply the end coordinate of the junction minus the start coordinate. By default, the junction must be between 70 bp and 500,000 bp, but the minimum and maximum can be set using “-i <minimum intron length>” and “-I <maximum intron length>”, respectively.

For efficiency, this tool can be used to process only alignments in a particular region as opposed to analyzing the entire BAM file. The option “-r <chr>:<start>-<stop>” can be used to set a single contiguous region of interest. Multiple jobs can be run in parallel to analyze separate non-contiguous regions.

#### Junctions annotate

This command takes a list of junctions in BED12 format as input and annotates them with respect to a reference transcriptome in GTF format. The observed splice-sites used are recorded based on a reference genome sequence in FASTA format. The output is written to STDOUT in TSV format, with separate columns for the number of splicing acceptors skipped, number of splicing donors skipped, number of exons skipped, the junction type, whether the donor site is known, whether the acceptor site is known, whether this junction is known, the overlapping transcripts, and the overlapping genes, in addition to the chromosome, start, stop, junction name, junction score, and strand taken from the input BED12 file. This output can be redirected to a file with “-o /PATH/TO/FILE”. By default, single exon transcripts are ignored in the GTF but can be included with the option “-S”.

#### Cis-splice-effects identify

This command combines the above utilities into a pipeline for identifying variants that may cause aberrant splicing events by altering splicing motifs in *cis*. As such, it relies on essentially the same inputs: a gzipped and Tabix-indexed VCF file containing a list of variants, an alignment BAM/CRAM file containing aligned RNA-seq reads, a GTF file containing the reference transcriptome of interest, and a FASTA file containing the reference genome sequence of interest.

First, the list of variants is annotated. The splice variant window size is set using the options “-e”, “-i”, “-E”, and “-I”, just as in *variants annotate*. The splice junction region size (i.e. the range around a particular variant in which an overlapping junction is associated with the variant) can be set using “-w <splice junction region size>”. By default, this range is not a particular number of bases but is calculated individually for each variant, depending on the variant type annotation. For “splicing_exonic”, “splicing_intronic”, and “exonic” variants, the region extends from the 3’ end of the exon directly upstream of the variant-associated exon to the 5’ end of the exon directly downstream of it. For “intronic” variants, the region is limited to the intron containing the variant. Single-exons can be kept with the “-S” option. The annotated list of variants in VCF format (analogous to the output of *variants annotate*) can be written to a file with “-v /PATH/TO/FILE”.

The BAM file is then processed based on the splice junction regions to produce the list of junctions present within these regions. A file containing these junctions in BED12 format (analogous to the output of *junctions extract*) can be written using “-j /PATH/TO/FILE”. The minimum anchor length, minimum intron length, and maximum intron length can be set using a”, “-i”, and “-I” options, just as in *junctions extract*.

The list of junctions produced by the preceding step is then annotated with the information presented in the *junctions annotate* section above. Additionally, each junction is annotated with a list of associated variants (i.e. variants whose splice junction regions overlapped the junction). The final output is written to STDOUT in TSV format (analogous to the output of *junctions annotate*) or can be redirected to a file with “-o /PATH/TO/FILE”.

#### Cis-splice-effects associate

This command is similar to *cis-splice-effects identify*, but takes the BED output of *junctions extract* in lieu of an alignment file with RNA alignments. As with *cis-splice-effects identify*, each junction is annotated with a list of associated variants (i.e. variants whose splice junction regions overlapped the junction). The resulting output is then the same as *cis-splice-effects identify*, but limited to the junctions provided as input.

### Analysis

#### Dataset Description

32 cancer cohorts were analyzed from TCGA. These cancer types are Adrenocortical carcinoma (ACC), Bladder Urothelial Carcinoma (BLCA), Brain Lower Grade Glioma (LGG), Breast invasive carcinoma (BRCA), Cervical squamous cell carcinoma and endocervical adenocarcinoma (CESC), Cholangiocarcinoma (CHOL), Colon adenocarcinoma (COAD), Esophageal carcinoma (ESCA), Glioblastoma multiforme (GBM), Head and Neck squamous cell carcinoma (HNSC), Kidney Chromophobe (KICH), Kidney renal clear cell carcinoma (KIRC), Kidney renal papillary cell carcinoma (KIRP), Liver hepatocellular carcinoma (LIHC), Lung adenocarcinoma (LUAD), Lung squamous cell carcinoma (LUSC), Lymphoid Neoplasm Diffuse Large B cell Lymphoma (DLBC), Mesothelioma (MESO), Ovarian serous cystadenocarcinoma (OV), Pancreatic adenocarcinoma (PAAD), Pheochromocytoma and Paraganglioma (PCPG), Prostate adenocarcinoma (PRAD), Rectum adenocarcinoma (READ), Sarcoma (SARC), Skin Cutaneous Melanoma (SKCM), Stomach adenocarcinoma (STAD), Testicular Germ Cell Tumors (TGCT), Thymoma (THYM), Thyroid carcinoma (THCA), Uterine Carcinosarcoma (UCS), Uterine Corpus Endometrial Carcinoma (UCEC), and Uveal Melanoma (UVM). Three cohorts were derived from patients at Washington University in St. Louis. These cohorts are Hepatocellular Carcinoma (HCC), Oral Squamous Cell Carcinoma (OSCC), and Small Cell Lung Cancer (SCLC).

#### Sample processing for cohorts with bulk transcriptome data

We applied RegTools to 35 tumor cohorts. Genomic and transcriptomic data for 32 cohorts were obtained from The Cancer Genome Atlas (TCGA). Information regarding the alignment and variant calling for these samples is described by the Genomic Data Commons data harmonization effort^72^. Whole exome sequencing (WES) mutation calls for these samples from MuSE^73^, MuTect2^74^, VarScan2^75^, and SomaticSniper^76^, were left-aligned, trimmed, and decomposed to ensure the correct representation of the variants across the multiple callers. Samples for the remaining three cohorts, HCC^77^, SCLC^34^, and OSCC^78^, were sequenced at Washington University in St. Louis. Genomic data were produced by WES for SCLC and OSCC and whole genome sequencing (WGS) for HCC. Normal genomic data of the same sequencing type and tumor RNA-seq data were also available for all subjects. Sequence data were aligned using the Genome Modeling System (GMS)^79^ using TopHat2 for RNA and BWA-MEM^80^ for DNA. HCC and SCLC were aligned to GRCh37 while OSCC was aligned to GRCh38. Somatic variant calls were made using Samtools v0.1.1^26^, SomaticSniper2 v1.0.2^76^, Strelka V0.4.6.2^81^, and VarScan v2.2.6^75,81^ through the GMS. High-quality mutations for all samples were then selected by requiring that a variant be called by two of the four variant callers.

Additional samples from orthogonal projects at Washington University in St. Louis were used for the orthogonal validation analysis. Samples included in this analysis were of the following cancer types: SCLC, GBM, and brain metastases that resulted from lung and breast cancers. The SCLC samples were an extension of the previous SCLC cohort we used for the pan-cancer analysis, with the difference being that the additional samples being aligned to GRCh38. Methods for the sample processing of the GBM and brain metastases samples are described in the original analysis manuscript for these data^33^.

#### Mice used for MCB6C experiments

5- to 6-week-old black 6 (B6NTac) male mice were purchased from Taconic Biosciences and were housed in a SPF barrier facility under the guidelines of Institutional Animal Care and Use Committee at Washington University. All the in vivo experiments were performed one week after mice were delivered to the animal facility.

#### Mouse bladder organoid culture for mouse injection

One previously archived frozen vial of singly suspended MCB6C organoid was thawed at least 2 weeks before mouse injection and expanded weekly in culture at least 2 times. For MCB6C organoid culture expansion, growth factor reduced Matrigel was thawed on ice for minimally 1 ½ hours. Pelleted MCB6C cells were washed and resuspended in 1 ml of Advanced DMEM/F12+++ medium (Advanced DMEM/F12 medium [125634028, Gibco] supplemented with 1% penicillin/streptomycin, 1% HEPEs and Glutamax) and cell concentration was determined by automated cell counter. To establish organoid culture, 50 ul Matrigel tabs with 10,000 cells/tab were generated and plated on 6-well suspension culture plates, 6 tabs wells. Tabs were incubated at 37C for 15 minutes until Matrigel was hardened, returned to tissue culture incubator, and cultured with mouse bladder organoid medium (MBO medium - Advanced DMEM/F12+++ medium supplemented with EGF, A-83-01, Noggin, R-Spondin, N-Acetly-L-cysteine and Nicotinamide). Organoids were replenished with fresh MBO medium every 3-4 days and also one day before mouse injection.

#### Mouse injection with MCB6C organoid cells

A single cell suspension of MCB6C organoid was generated by TrypLE Express(12605010, Gibco) digestion organoid Matrigel tabs at 37C for 15 minutes. After digestion, pelleted cells were washed and resuspended in PBS to determine cell concentration. After cell concentration was adjusted to 20 million/ml in PBS, organoid cells were mixed with growth factor reduced Matrigel at 1:1 ratio before injected subcutaneously into the left flank of the mouse (1 million/100ul cells each mouse). Tumor development was monitored using digital calipers to assess the length, width, and depth of each tumor. For ICB, each mouse was injected intraperitoneally with 250 ug anti-PD1 (RMP1-14, BioXcell) and 200 ug anti-CTLA-4 (9D9, BioXcell) day 9 and 12 after organoid implantation. For isotype controls, each mouse was injected with 250 ug rat IgG2a (2A3, BioXcell) and 200 ug IgG2b (MPC-11, BioXcell). For CD4+ T cell depletion, each mouse was injected with 250 ug anti-CD4 (GK1.5, BioXcell) day 0 and 7 after organoid depletion. Rat IgG2b (LTF-2, BioXcell) was used as isotype control for anti-CD4.

#### Harvesting MCB6C tumors for single cell RNA-seq analysis

Based on 10x Genomics Demonstrated Protocols, 14 days after organoid implantation, tumors were dissected from euthanized mice, cut into small pieces of ~2-4 mm^3^, and further processed into dead-cell depleted single cell suspension following manufacturer’s protocol using Tumor Dissociation Kit and MACS Dead Cell removal Kit (Miltenyi Biotec). Briefly, tumor tissue pieces were transferred to gentleMACS C tube containing enzyme mix before loading onto a gentleMACS Octo Dissociator with Heaters for tissue digestion at 37C for 80 minutes. After tissue dissociation was completed, cell suspension was transferred to a new 50 ml conical tube, and supernatant was removed after centrifugation. Cell pellet was resuspended in RPMI 1640 medium, filtered through a prewetted 70 uM cell filter, strained, pelleted, and resuspended in red cell lysis buffer and incubated on ice for 10 minutes. After adding the wash buffer, the cell suspension was pelted and resuspended in the wash buffer. To remove dead cells, Dead Cell Removal Microbeads were added to resuspend cell pellet (100 ul beads per 10^7 cells) using a wide-bore pipette tip. After incubation for 15 minutes at room temperature, the cell-microbead mixture was applied onto a MS column. Dead cells remained in the column and the effluent represented to the live cell fraction. The percentage of viable cells was determined by an automated cell counter. Dead cell removal was repeated if the percentage of viable cells did not reach above 90%. Two rounds of centrifugation/resuspension were carefully performed for two rounds in 1xPBS/0.04% BSA using a wide-bore tip. To submit cell samples for single-cell RNA-seq analysis, cell concentration was determined accurately by sampling cell suspension twice and counting each sampling twice and adjusted to 1,167 cells/ul.

#### Single-cell RNA-seq analysis of MCB6C cells

40 ul of each cell suspension was submitted to the Genome Technology Access Center/McDonnell Genome Institute (GTAC/MGI) for 10x Genomics single cell RNA-seq analysis using the 5’v2 library kit of TotalSeq C antibodies with BCR and TCR V(D)J enrichment. FASTQs and Cell Ranger output was generated. Alignment and gene expression quantification done using CellRanger count (v5.0). Matrices are then imported into Seurat^82^ (v4.0.1) for filtering cells, QC, clustering, etc. To filter suspected dying cells, cells were clustered before filtering and cells clustering based on high mitochondrial gene expression were identified. The cutoff of mitochondrial expression was based on the expression level that captures most of these cells. Doublets were filtered based on high UMI expression, with the top 0.9% of genes removed from each condition in each replicate. Cutoffs for filtering of cells with low feature detection was done by assigning cell type to each cell with CellMatch, identifying cells that did not have enough features for their cell type to be predicted, and identifying average feature expression in these cells. Aftering filtered cells were removed, remaining cells were scaled, normalized, and clustered following Seurat’s vignette.

#### Long read sequencing of HCC1395 cell line

The HCC1395 cell line is described as being of tissue origin: mammary gland; breast/duct. The patient’s cancer was described as: TNM stage I, grade 3, primary ductal carcinoma. The patient received chemotherapy prior to isolation of the tumor^83^. This tumor is considered “Triple-Negative” by classic typing: ERBB2-negative (aka HER2/neu), PR-negative, and ER-negative. Otherwise, it is one of those difficult to classify by expression-based molecular typing but is likely of the “Basal” sub-type ^84^. For a normal comparator, we used HCC1395’s matched lymphoblastoid cell line, HCC1395BL. The HCC1395BL cell line was created from a B lymphoblast that was transformed by the EBV virus. For each of these samples, WGS, WES, and bulk RNA-seq were performed. Whole-genome sequencing was performed to a target median coverage depth of ~30x for the normal samples and ~50x for the tumor sample. Exome sequencing was performed to a target median coverage depth of ~100x. RNA-seq was performed for both tumor and normal. Additionally, Oxford Nanopore Technologies long-read sequencing was performed using both the Direct RNA Sequencing Kit and Direct cDNA Sequencing Kit. The Direct RNA Sequencing Kit yielded 1.1 million reads with 1.07 Gb of passed bases and read lengths ranging from ~500 basepairs (bp) to ~8 kilobases (kb). The Direct cDNA Sequencing Kit was run twice. The first run used the RNA from the same mRNA enrichment as the RNA Direct library. This sequencing run yielded 2.48 million reads with 2.36 Gb of passed bases and read lengths ranging from ~500 bp to ~9.6 kb. The second Direct cDNA Sequencing Kit was applied to a new RNA extraction, so a separate mRNA enrichment from the first two runs. This run yielded 6.6 million reads with 4.05 Gb passed bases and read lengths ranging from ~500 basepairs (bp) to ~8 kilobases (kb). These data were aligned to GRCh38 using recommended settings for minimap2^71^. To confirm junctions identified by Illumina sequencing, junctions were extracted from ONT alignment files and combined for the three libraries. For each junction we were attempting to validate, we required that there be at least one read of support that utilized either the donor or the acceptor site of the junction of interest. If there was no evidence of either donor or acceptor site being used, we concluded that we had insufficient coverage to validate that particular junction.

#### Candidate junction filtering

To generate results for 4 splice variant window sizes, we ran *cis-splice-effects identify* with 4 sets of splice variant window parameters. For our “i2e3” window (RegTools default), to examine intronic variants within 2 bases and exonic variants within 3 bases of the exon edge, we set “-i 2 -e 3”. Similarly, for “i50e5”, to examine intronic variants within 50 bases and exonic variants within 5 bases of the exon edge, we set “-i 50 -e 5”. To view all exonic variants, we simply set “-E”, without “-i” or “-e” options. To view all intronic variants, we simply set “-I”, without “-i” or “-e” options. TCGA samples were processed with GRCh38.d1.vd1.fa (downloaded from the GDC reference file page at https://gdc.cancer.gov/about-data/gdc-data-processing/gdc-reference-files) as the reference fasta file and gencode.v29.annotation.gtf (downloaded via the GENCODE FTP) as the reference transcriptome. OSCC was processed with Homo_sapiens.GRCh38.dna_sm.primary_assembly.fa and Homo_sapiens.GRCh38.79.gtf (both downloaded from Ensembl). HCC and SCLC were processed with Homo_sapiens.GRCh37.dna_sm.primary_assembly.fa and Homo_sapiens.GRCh37.87.gtf (both downloaded from Ensembl).

#### Statistical filtering of candidate events

We refer to a statistical association between a variant and a junction as an “event”. For each event identified by RegTools, a normalized score (norm_score) was calculated for the junction of the event by dividing the number of reads supporting that junction by the sum of all reads for all junctions within the splice junction region for the variant of interest. This metric is conceptually similar to a “percent-spliced in” (PSI) index, but measures the presence of entire exon-exon junctions, instead of just the inclusion of individual exons. If there were multiple samples that contained the variant for the event, then the mean of the normalized scores for the samples was computed (mean_norm_score). If only one sample contained the variant, its mean_norm_score is equal to its norm_score. This value was then compared to the distribution of samples which did not contain the variant to calculate a p-value as the percentage of the norm_scores from these samples which are at least as high as the mean_norm_score computed for the variant-containing samples. We performed separate analyses for events involving known junctions (DA) and those involving novel junctions which used at least one known splice site (D/A/NDA), based on annotations in the corresponding reference GTF. For this study, we filtered out any junctions which did not use at least one known splice site (N) and junctions which did not have at least 5 reads of evidence across variant-containing samples. The Benjamini-Hochberg procedure was then applied to the remaining events. Following correction, an event was considered significant if its adjusted p-value was ≤ 0.05.

#### Annotation with GTEx junction data and other splice prediction tools

Events identified by RegTools as significant were annotated with information from GTEx, VEP, CTAT splicing, SpliceAI, MiSplice, and Veridical. GTEx junction information was obtained from the GTEx Portal. Specifically, the exon-exon junction read counts file from the v8 release was used for data aligned to GRCh38 while the same file from the v7 release was used for the data aligned to GRCh37. Mappings between tumor cohorts and GTEx tissues can be found in **Supplementary File 15**. We annotated all starting variants with VEP in the “per_gene” and “pick” modes. The “per_gene” setting outputs only the most severe consequence per gene while the “pick” setting picks one line or block of consequence data per variant. We considered any variant with at least one splice-associated annotation to be “VEP significant”. All variants were also processed with SpliceAI using the default options. A variant was considered to be “SpliceAI significant” if it had at least one score greater than 0.2, the developers’ recommended value for high recall of their model. Instructions and scripts to annotate with GTEx and SpliceAI are available at regtools.org and in the RegTools GitHub repository. Variants identified by MiSplice^20^ were obtained from the paper’s supplemental tables and were lifted over to GRCh38. Variants identified by SAVNet^23^ were obtained from the paper’s supplemental tables and were lifted over to GRCh38. Variants identified by Veridical^21,22^ were obtained via download from the link referenced within the manuscript and lifted over to GRCh38.

#### Visual exploration of statistically significant candidate events

IGV sessions were created for each event identified by RegTools that was statistically significant. Each IGV session file contained a bed file with the junction, a vcf file with the variant, and an alignment file for each sample that contained the variant. Additional information, such as the splice sites predicted by SpliceAI, were also added to these session files to enhance the exploration of these events. Events of interest were manually reviewed in IGV to assess whether the association between the variant and junction made sense in a biological context (e.g. affected a known splice site, altered a genomic sequence to look more like a canonical splice site, or the novel junction disrupted active or regulatory domains of the protein product). An extensive review of literature and visualizations of junction usage in the presence and absence of the variant were also used to identify novel, biologically relevant events.

#### Identification of genes with recurrent splice-associated variants

For each cohort, we calculated a p-value to assess whether the splicing profile from a particular gene was significantly more likely to be altered by somatic variants. Specifically, we performed a 1-tailed binomial test, considering the number of samples in a cohort as the number of attempts. Success was defined by whether the sample had evidence of at least one splice-associated variant in that gene. The null probability of success, *p*_null_ was calculated as

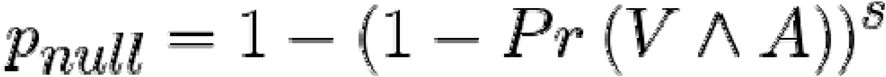

where *s* is the total number of base positions residing in any of the gene’s splice variant windows, *V* is the event that a somatic variant occurred at such a base position, and *A* is the event that this variant was deemed to be significantly associated with at least one junction in our analysis. The joint probability that both V and A occurred was estimated by dividing the total of events across all samples in which each junction was detected by *s*. The value of *s* was computed based on the exon and transcript definitions in the reference GTF used for performing RegTools analyses on a given cohort.

We also calculated overall metrics, in order to rank genes. For each set of cohorts (e.g. TCGA-only, MGI-only, combined), an overall p-value was computed for each gene according to the above formula, pooling all of the samples across the included cohorts, and the fraction of samples was simply calculated by dividing the number of samples in which an event occurred within the given gene by the total number of samples, pooled across the included cohorts. The reference GTF used for analyzing the TCGA samples (i.e. gencode.v29.annotation.gtf) was used for all sets of cohorts.

## Code availability

RegTools is open source (MIT license) and available at https://github.com/griffithlab/regtools/. All scripts used in the analyses presented here are also provided. For ease of use, a Docker container has been created with RegTools, SpliceAI, R, and Python 3 installed (https://hub.docker.com/r/griffithlab/regtools/). This Docker container allows a user to run the workflow we outline at https://regtools.readthedocs.io/en/latest/workflow/. Docker is an opensource software platform that enables applications to be readily installed and run on any system. The availability of RegTools with all its dependencies as a Docker container also facilitates the integration of the RegTools software into workflow pipelines that support Docker images.

## Data availability

Sequence data for each cohort analyzed in this study are available through dbGaP at the following accession IDs: phs000178 for TCGA cohorts, phs001106 for HCC, phs001049 for SCLC, phs001623 for OSCC, and phs002612.v1.p1 for GBM/Brain metastases. Statistically significant events for D, A, and NDA junctions across the four variant splicing windows used are available via **Supplementary Files 1 and 2**. Statistically significant events for DA junctions are available as **Supplementary Files 3 and 4**. Complete results of gene recurrence analysis are available as **Supplementary Files 10-14**.

## Acknowledgments

We thank the patients and their families for the donation of their samples and participation in clinical trials. We would like to thank Donald Conrad for his initial idea to compare to variant effect predictor tools. Kelsy Cotto was supported by Siteman Cancer Center under fund number #3477-92400 and T32CA113275. Avinash Ramu was supported by the Burroughs Wellcome Fund Institutional Program Unifying Population and Laboratory Based Sciences Award at Washington University. Malachi Griffith was supported by the National Human Genome Research Institute (NHGRI) of the National Institutes of Health (NIH) under Award Number R00HG007940. Malachi Griffith and Obi Griffith were supported by the NIH National Cancer Institute (NCI) under Award Numbers U01CA209936, U01CA231844, U01CA248235 U24CA237719. Malachi Griffith and Megan Richters were supported by the V Foundation for Cancer Research under Award Number V2018-007. The results published here are in whole or part based upon data generated by the TCGA Research Network: https://www.cancer.gov/tcga.

## Contributions

K.C.C. and Y.-Y.F. were involved in all aspects of this study, including designing methodology, developing and testing the tool software, analyzing and interpreting data, and writing the manuscript, with input from A.R., M.R., S.L.F., Z.L.S., H.X., J.F.M., J.K., K.M.C., T.H.C., E.B.R., D.A., S.D., O.L.G., and M.G. A.R. designed the tool and led software development efforts. Y.L., W.C.C., C.A.M., V.A., G.P.D., R.U., and R.G. provided unpublished tumor datasets and provided critical feedback on the manuscript. O.L.G. and M.G. supervised the study. All authors read and approved the final manuscript.

## References

1. Chabot, B. & Shkreta, L. Defective control of pre-messenger RNA splicing in human disease. J. Cell Biol. 212, 13–27 (2016).

2. Vogelstein, B. et al. Cancer genome landscapes. Science 339, 1546–1558 (2013).

3. Soemedi, R. et al. Pathogenic variants that alter protein code often disrupt splicing. Nat. Genet. 49, 848–855 (2017).

4. Supek, F., Miñana, B., Valcárcel, J., Gabaldón, T. & Lehner, B. Synonymous mutations frequently act as driver mutations in human cancers. Cell 156, 1324–1335 (2014).

5. Jung, H. et al. Intron retention is a widespread mechanism of tumor-suppressor inactivation. Nat. Genet. 47, 1242–1248 (2015).

6. Venables, J. P. Aberrant and alternative splicing in cancer. Cancer Res. 64, 7647–7654 (2004).

7. Climente-González, H., Porta-Pardo, E., Godzik, A. & Eyras, E. The Functional Impact of Alternative Splicing in Cancer. Cell Rep. 20, 2215–2226 (2017).

8. Chen, J. & Weiss, W. A. Alternative splicing in cancer: implications for biology and therapy. Oncogene 34, 1–14 (2015).

9. Xiong, H. Y. et al. RNA splicing. The human splicing code reveals new insights into the genetic determinants of disease. Science 347, 1254806 (2015).

10. Yeo, G. & Burge, C. B. Maximum entropy modeling of short sequence motifs with applications to RNA splicing signals. J. Comput. Biol. 11, 377–394 (2004).

11. Fairbrother, W. G., Yeh, R.-F., Sharp, P. A. & Burge, C. B. Predictive identification of exonic splicing enhancers in human genes. Science 297, 1007–1013 (2002).

12. Wang, Z. et al. Systematic identification and analysis of exonic splicing silencers. Cell 119, 831–845 (2004).

13. Jaganathan, K. et al. Predicting Splicing from Primary Sequence with Deep Learning. Cell 176, 535–548.e24 (2019).

14. Kahles, A., Ong, C. S., Zhong, Y. & Rätsch, G. SplAdder: identification, quantification and testing of alternative splicing events from RNA-Seq data. Bioinformatics 32, 1840–1847 (2016).

15. Trincado, J. L. et al. SUPPA2: fast, accurate, and uncertainty-aware differential splicing analysis across multiple conditions. Genome Biol. 19, 40 (2018).

16. Kahles, A. et al. Comprehensive Analysis of Alternative Splicing Across Tumors from 8,705 Patients. Cancer Cell 34, 211–224.e6 (2018).

17. Li, Y. I. et al. Annotation-free quantification of RNA splicing using LeafCutter. Nat. Genet. 50, 151–158 (2018).

18. Monlong, J., Calvo, M., Ferreira, P. G. & Guigó, R. Identification of genetic variants associated with alternative splicing using sQTLseekeR. Nat. Commun. 5, 4698 (2014).

19. Li, Y. I. et al. RNA splicing is a primary link between genetic variation and disease. Science 352, 600–604 (2016).

20. Jayasinghe, R. G. et al. Systematic Analysis of Splice-Site-Creating Mutations in Cancer. Cell Rep. 23, 270–281.e3 (2018).

21. Viner, C., Dorman, S. N., Shirley, B. C. & Rogan, P. K. Validation of predicted mRNA splicing mutations using high-throughput transcriptome data. F1000Res. 3, (2014).

22. Shirley, B. C., Mucaki, E. J. & Rogan, P. K. Pan-cancer repository of validated natural and cryptic mRNA splicing mutations. F1000Res. 7, 1908 (2018).

23. Shiraishi, Y. et al. A comprehensive characterization of cis-acting splicing-associated variants in human cancer. Genome Res. 28, 1111–1125 (2018).

24. GTEx Consortium. The Genotype-Tissue Expression (GTEx) project. Nat. Genet. 45, 580–585 (2013).

25. McLaren, W. et al. The Ensembl Variant Effect Predictor. Genome Biol. 17, 122 (2016).

26. Li, H. et al. The Sequence Alignment/Map format and SAMtools. Bioinformatics 25, 2078–2079 (2009).

27. Sondka, Z. et al. The COSMIC Cancer Gene Census: describing genetic dysfunction across all human cancers. Nat. Rev. Cancer 18, 696–705 (2018).

28. Robinson, J. T. et al. Integrative genomics viewer. Nat. Biotechnol. 29, 24–26 (2011).

29. Palmisano, A., Vural, S., Zhao, Y. & Sonkin, D. MutSpliceDB: A database of splice sites variants with RNA-seq based evidence on effects on splicing. Hum. Mutat. 42, 342–345 (2021).

30. Barretina, J. et al. The Cancer Cell Line Encyclopedia enables predictive modelling of anticancer drug sensitivity. Nature 483, 603–607 (2012).

31. Chang, L.-C., Vural, S. & Sonkin, D. Detection of homozygous deletions in tumorsuppressor genes ranging from dozen to hundreds nucleotides in cancer models. Hum. Mutat. 38, 1449–1453 (2017).

32. Ghandi, M. et al. Next-generation characterization of the Cancer Cell Line Encyclopedia. Nature 569, 503–508 (2019).

33. Schaettler, M. O. et al. Characterization of the Genomic and Immunologic Diversity of Malignant Brain Tumors through Multisector Analysis. Cancer discovery vol. 12 154–171 (2022).

34. Wagner, A. H. et al. Recurrent WNT pathway alterations are frequent in relapsed small cell lung cancer. Nat. Commun. 9, 3787 (2018).

35. Sato, Y. et al. CD4+ T cells induce rejection of urothelial tumors after immune checkpoint blockade. JCI Insight 3, (2018).

36. UniProt Consortium. UniProt: the universal protein knowledgebase in 2021. Nucleic Acids Res. 49, D480–D489 (2021).

37. Rui, Y. et al. Axin stimulates p53 functions by activation of HIPK2 kinase through multimeric complex formation. EMBO J. 23, 4583–4594 (2004).

38. Lundgaard, G. L. et al. Identification of a novel effector domain of BIN1 for cancer suppression. J. Cell. Biochem. 112, 2992–3001 (2011).

39. Ghaneie, A. et al. Bin1 attenuation in breast cancer is correlated to nodal metastasis and reduced survival. Cancer Biol. Ther. 6, 192–194 (2007).

40. Zhong, X. et al. Bin1 is linked to metastatic potential and chemosensitivity in neuroblastoma. Pediatr. Blood Cancer 53, 332–337 (2009).

41. Gurumurthy, S., Vasudevan, K. M. & Rangnekar, V. M. Regulation of apoptosis in prostate cancer. Cancer Metastasis Rev. 20, 225–243 (2001).

42. Xie, X., Zheng, X., Xie, T., Cai, J. & Yao, Y. Identification of prognostic alternative splicing signatures in uveal melanoma. Int. Ophthalmol. 41, 1347–1362 (2021).

43. Surget, S., Khoury, M. P. & Bourdon, J.-C. Uncovering the role of p53 splice variants in human malignancy: a clinical perspective. Onco. Targets. Ther. 7, 57–68 (2013).

44. Tokheim, C. & Karchin, R. CHASMplus Reveals the Scope of Somatic Missense Mutations Driving Human Cancers. Cell Syst 9, 9–23.e8 (2019).

45. Cui, M. et al. Immunoglobulin Expression in Cancer Cells and Its Critical Roles in Tumorigenesis. Front. Immunol. 12, 613530 (2021).

46. Chu, J. et al. IGHG1 Regulates Prostate Cancer Growth via the MEK/ERK/c-Myc Pathway. Biomed Res. Int. 2019, 7201562 (2019).

47. Li, Y. et al. IGHG1 induces EMT in gastric cancer cells by regulating TGF-β/SMAD3 signaling pathway. J. Cancer 12, 3458–3467 (2021).

48. Li, X. et al. IGHG1 upregulation promoted gastric cancer malignancy via AKT/GSK-3β/β-Catenin pathway. Cancer Cell Int. 21, 397 (2021).

49. Bonneville, R. et al. Landscape of Microsatellite Instability Across 39 Cancer Types. JCO Precis Oncol 2017, (2017).

50. Kloor, M. et al. Immunoselective pressure and human leukocyte antigen class I antigen machinery defects in microsatellite unstable colorectal cancers. Cancer Res. 65, 6418–6424 (2005).

51. Sade-Feldman, M. et al. Resistance to checkpoint blockade therapy through inactivation of antigen presentation. Nat. Commun. 8, 1136 (2017).

52. Seliger, B., Maeurer, M. J. & Ferrone, S. Antigen-processing machinery breakdown and tumor growth. Immunol. Today 21, 455–464 (2000).

53. Güssow, D. et al. The human beta 2-microglobulin gene. Primary structure and definition of the transcriptional unit. J. Immunol. 139, 3132–3138 (1987).

54. Wang, L., Yin, W. & Shi, C. E3 ubiquitin ligase, RNF139, inhibits the progression of tongue cancer. BMC Cancer 17, 452 (2017).

55. Hornbeck, P. V. et al. PhosphoSitePlus, 2014: mutations, PTMs and recalibrations. Nucleic Acids Res. 43, D512–20 (2015).

56. Zhao, R., Choi, B. Y., Lee, M.-H., Bode, A. M. & Dong, Z. Implications of Genetic and Epigenetic Alterations of CDKN2A (p16(INK4a)) in Cancer. EBioMedicine 8, 30–39 (2016).

57. Gump, J., Stokoe, D. & McCormick, F. Phosphorylation of p16 INK4A Correlates with Cdk4 Association. J. Biol. Chem. 278, 6619–6622 (2003).

58. Dobin, A. et al. STAR: ultrafast universal RNA-seq aligner. Bioinformatics 29, 15–21 (2013).

59. Kim, D., Langmead, B. & Salzberg, S. L. HISAT: a fast spliced aligner with low memory requirements. Nat. Methods 12, 357–360 (2015).

60. Kim, D. et al. TopHat2: accurate alignment of transcriptomes in the presence of insertions, deletions and gene fusions. Genome Biol. 14, R36 (2013).

61. Ellrott, K. et al. Scalable Open Science Approach for Mutation Calling of Tumor Exomes Using Multiple Genomic Pipelines. Cell Syst 6, 271–281.e7 (2018).

62. Takaku, M., Grimm, S. A. & Wade, P. A. GATA3 in Breast Cancer: Tumor Suppressor or Oncogene? Gene Expr. 16, 163–168 (2015).

63. Afzaljavan, F., Sadr, A. S., Savas, S. & Pasdar, A. GATA3 somatic mutations are associated with clinicopathological features and expression profile in TCGA breast cancer patients. Sci. Rep. 11, 1679 (2021).

64. Wang, Z. & Burge, C. B. Splicing regulation: from a parts list of regulatory elements to an integrated splicing code. RNA 14, 802–813 (2008).

65. Muro, A. F. et al. Regulation of fibronectin EDA exon alternative splicing: possible role of RNA secondary structure for enhancer display. Mol. Cell. Biol. 19, 2657–2671 (1999).

66. Schaal, T. D. & Maniatis, T. Multiple distinct splicing enhancers in the protein-coding sequences of a constitutively spliced pre-mRNA. Mol. Cell. Biol. 19, 261–273 (1999).

67. Black, D. L. A simple answer for a splicing conundrum. Proceedings of the National Academy of Sciences of the United States of America vol. 102 4927–4928 (2005).

68. Quinlan, A. R. BEDTools: The Swiss-Army Tool for Genome Feature Analysis. Curr. Protoc. Bioinformatics 47, 11.12.1–34 (2014).

69. Li, H. Tabix: fast retrieval of sequence features from generic TAB-delimited files. Bioinformatics 27, 718–719 (2011).

70. Bray, N. L., Pimentel, H., Melsted, P. & Pachter, L. Near-optimal probabilistic RNA-seq quantification. Nat. Biotechnol. 34, 525–527 (2016).

71. Li, H. Minimap2: pairwise alignment for nucleotide sequences. Bioinformatics 34, 3094–3100 (2018).

72. GDC Data Processing. https://gdc.cancer.gov/about-data/gdc-data-processing.

73. Fan, Y. et al. Accounting for tumor heterogeneity using a sample-specific error model improves sensitivity and specificity in mutation calling for sequencing data. bioRxiv 055467 (2016) doi:10.1101/055467.

74. Cibulskis, K. et al. Sensitive detection of somatic point mutations in impure and heterogeneous cancer samples. Nat. Biotechnol. 31, 213–219 (2013).

75. Koboldt, D. C. et al. VarScan 2: somatic mutation and copy number alteration discovery in cancer by exome sequencing. Genome Res. 22, 568–576 (2012).

76. Larson, D. E. et al. SomaticSniper: identification of somatic point mutations in whole genome sequencing data. Bioinformatics 28, 311–317 (2012).

77. Skidmore, Z. L. et al. Genomic and transcriptomic somatic alterations of hepatocellular carcinoma in non-cirrhotic livers. Cancer Genet. 264-265, 90–99 (2022).

78. Campbell, K. M. et al. Oral Cavity Squamous Cell Carcinoma Xenografts Retain Complex Genotypes and Intertumor Molecular Heterogeneity. Cell Rep. 24, 2167–2178 (2018).

79. Griffith, M. et al. Genome Modeling System: A Knowledge Management Platform for Genomics. PLoS Comput. Biol. 11, e1004274 (2015).

80. Li, H. & Durbin, R. Fast and accurate short read alignment with Burrows-Wheeler transform. Bioinformatics 25, 1754–1760 (2009).

81. Saunders, C. T. et al. Strelka: accurate somatic small-variant calling from sequenced tumor–normal sample pairs. Bioinformatics 28, 1811–1817 (2012).

82. Hao, Y. et al. Integrated analysis of multimodal single-cell data. Cell 184, 3573–3587.e29 (2021).

83. Gazdar, A. F. et al. Characterization of paired tumor and non-tumor cell lines established from patients with breast cancer. Int. J. Cancer 78, 766–774 (1998).

84. Heiser, L. M. et al. Subtype and pathway specific responses to anticancer compounds in breast cancer. Proc. Natl. Acad. Sci. U. S. A. 109, 2724–2729 (2012).

