## Supplemental Files/Figures for "RegTools: Integrated analysis of genomic and transcriptomic data for the discovery of splice-associated variants in cancer"

### Supplementary Files:

**Supplementary File 1**: Candidate variant junction pairings for D, A, NDA junctions (GRCh38 samples)

**Supplementary File 2**: Candidate variant junction pairings for D, A, NDA junctions (GRCh37 samples)

**Supplementary File 3**: Candidate variant junction pairings for DA junctions (GRCh38 samples)

**Supplementary File 4**: Candidate variant junction pairings for DA junctions (GRCh37 samples)

**Supplementary File 5**: Validation of MiSplice mini-gene assay findings using RegTools

**Supplementary File 6**: MutSpliceDB splice-associated variants validated using RegTools

**Supplementary File 7**: RegTools analysis for GBM/Brain metastases samples with multi-sector sequencing

**Supplementary File 8**: RegTools analysis for SCLC samples with treatment-naive and recurrence timepoints

**Supplementary File 9**: RegTools analysis of HCC1395 cell line with correlated Oxford Nanopore sequencing

**Supplementary File 10**: Recurrence analysis for D, A, NDA junctions in the default splice variant window

**Supplementary File 11**: Recurrence analysis for D, A, NDA junctions in the i50e5 splice variant window

**Supplementary File 12**: Recurrence analysis for D, A, NDA junctions in the E (all exonic) splice variant window

**Supplementary File 13**: Recurrence analysis for D, A, NDA junctions in the I (all intronic) splice variant window

**Supplementary File 14**: Recurrence analysis for DA junctions in the default splice variant window

**Supplementary File 15**: GTEx tissue type mapping to TCGA cohorts

### Supplemental Figures

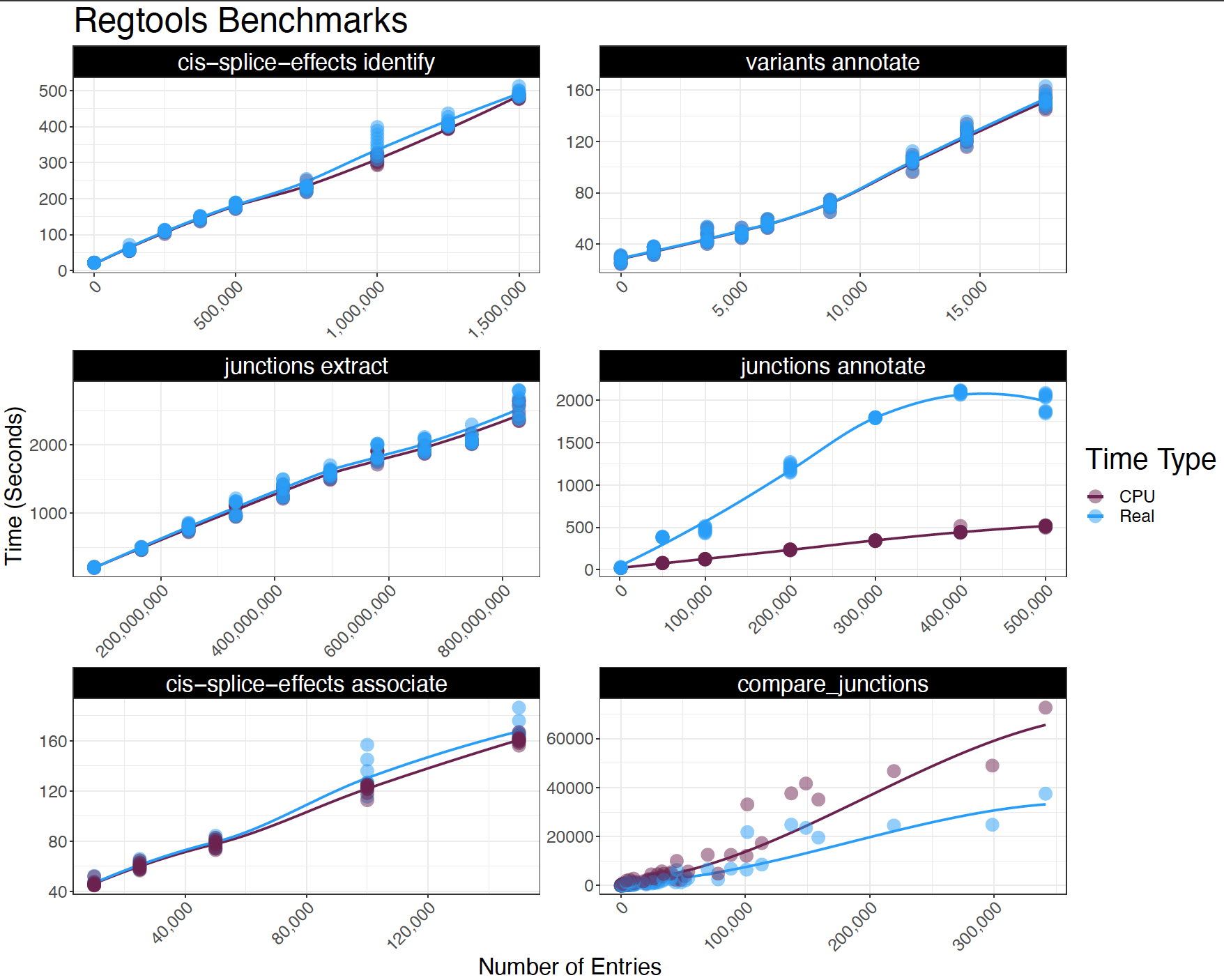

#### Supplementary Figure 1. Computational efficiency of Regtools

The total CPU time (System Time + User Time) and real-time are plotted against the number of entries processed for each available RegTools function using ten replicates. For the cis-splice-effects identify/cis-splice-effects associate/variants annotate workflows, the number of entries corresponds to the number of somatic variants. In contrast, the number of entries in the junctions extract/junctions annotate/compare_junctions workflows corresponds to the number of reads processed from a downsampled BAM file, the number of junctions processed, and the number of candidate variant junction pairings processed, respectively. For the statistical filtering performed by the supplementary compare_junctions pipeline, candidate variant junction pairings were compared across the number of samples in that cohort, with the largest being 1,022 samples that comprise our BRCA cohort. LOESS curves are fitted onto each plot.

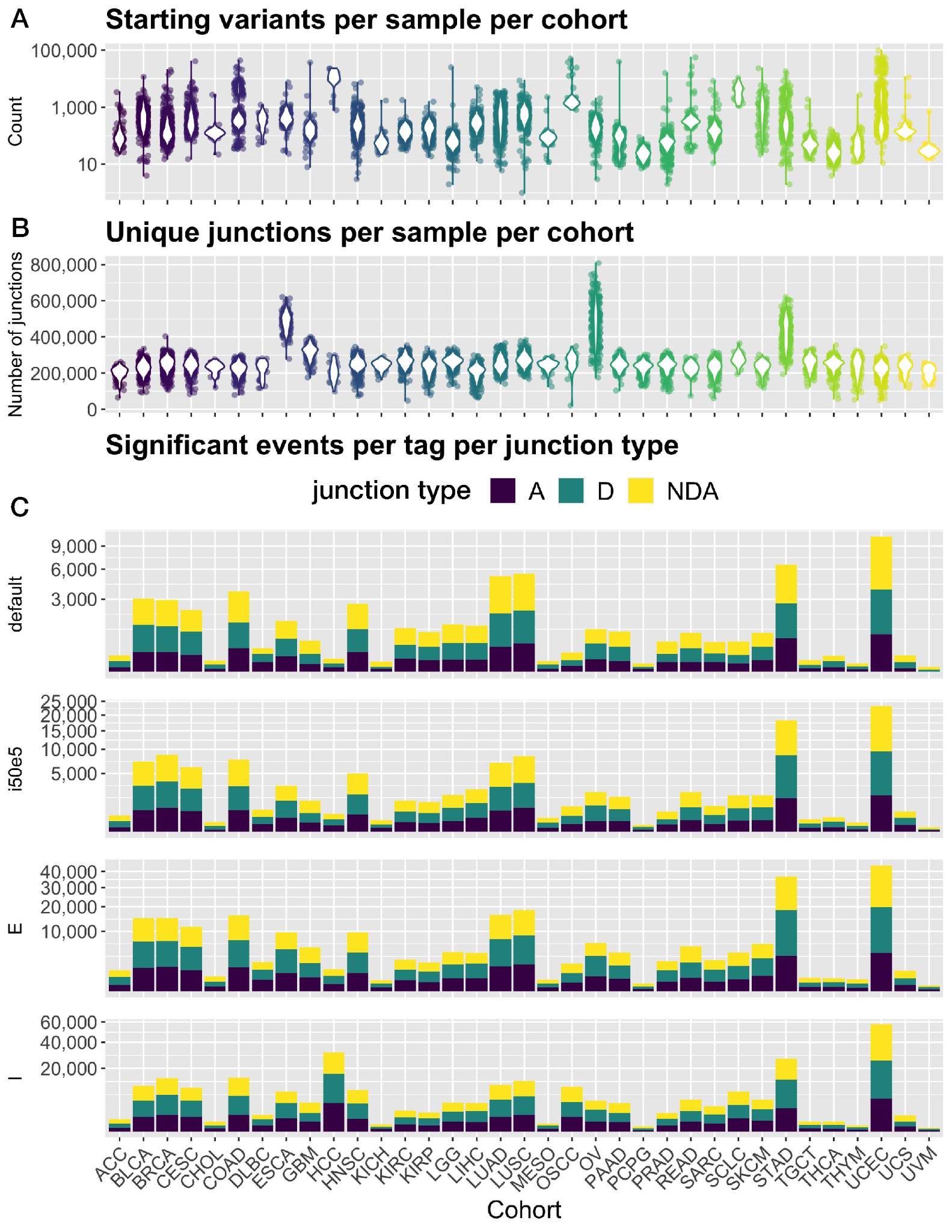

#### Supplementary Figure 2. Overview of input data considered and significant events identified by RegTools for each tumor type.

A) Summary of initial variants considered for analysis by RegTools per sample per tumor cohort. Each sample’s variant count is plotted, and violin plots are overlaid for each cohort. B) Summary of unique exon-exon junction observations for each sample. Each sample’s unique junction count is plotted, and violin plots are overlaid for each cohort. C) Summary of significant junction types for each cohort for each of the splice variant window parameters that were used in this analysis.

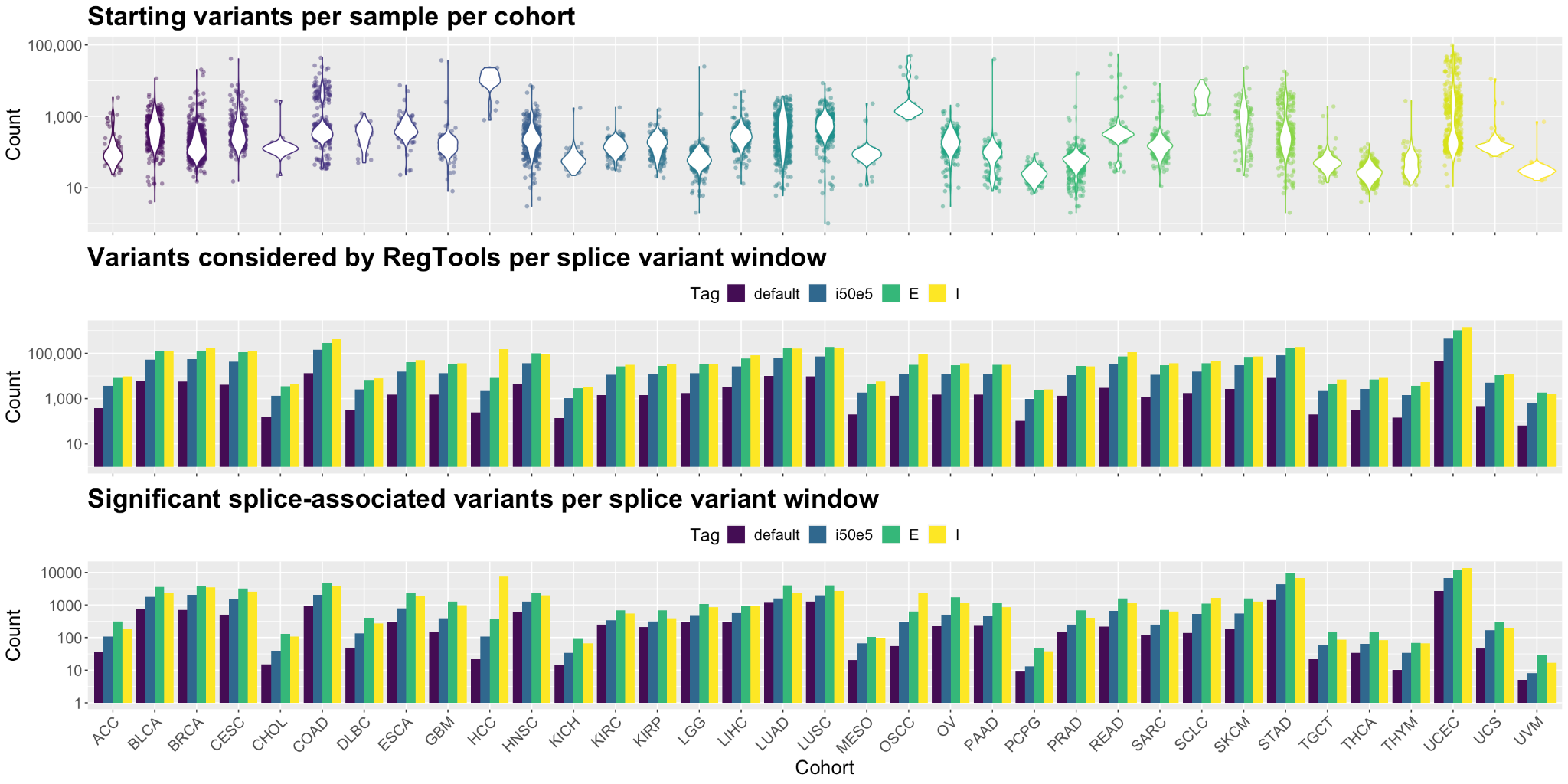

#### Supplementary Figure 3. Summary of variants analyzed by RegTools in each tumor cohort

Summary of the starting number of high-quality variants per sample, the number of initial variants considered for analysis by RegTools for each variant window used per tumor cohort, and the number of significant variants for each variant window used per tumor cohort.

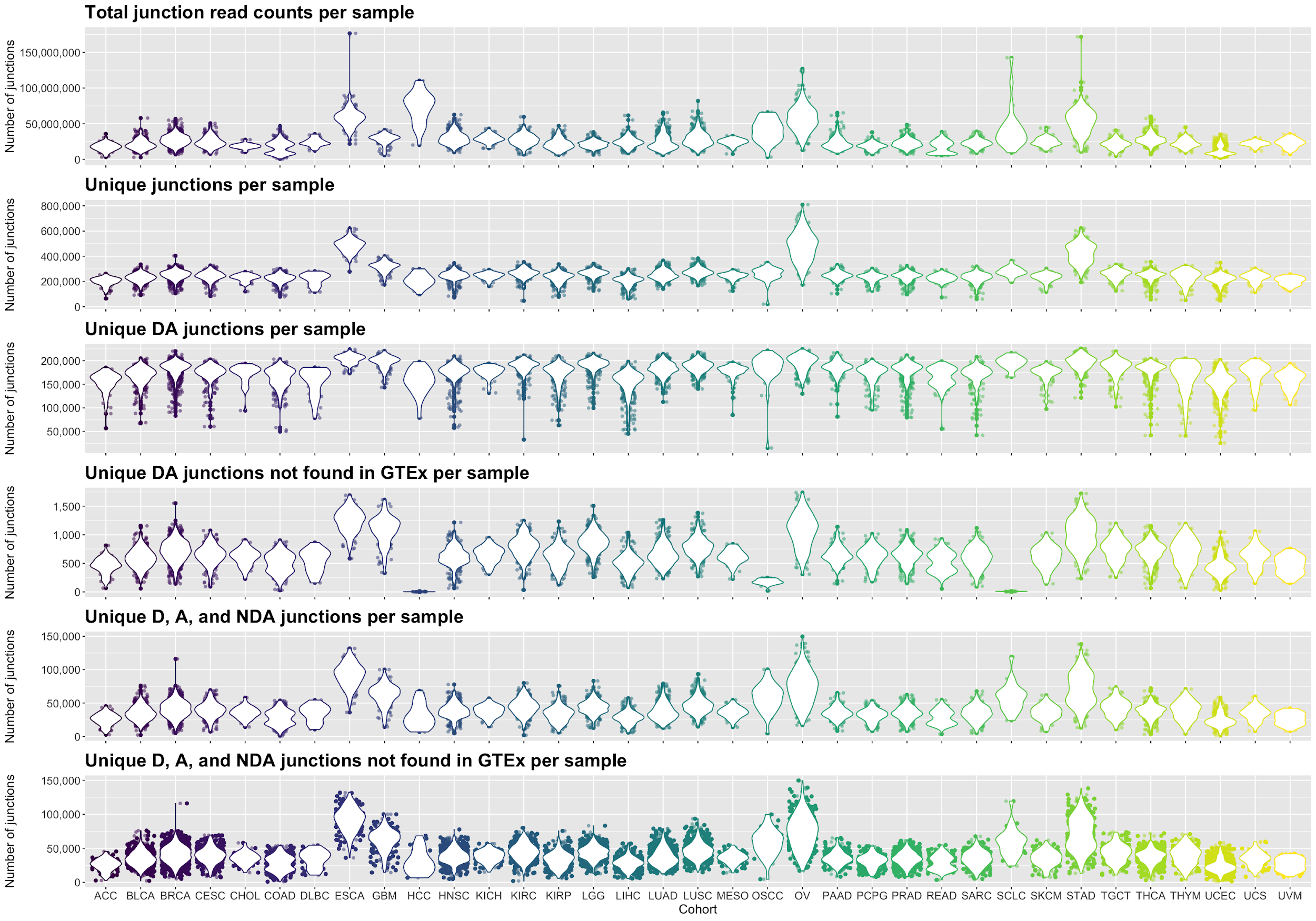

#### Supplementary Figure 4. Diversity of alternative splicing patterns across cancer types.

Summary of the total junction read counts, unique junctions (all types), unique known (DA) junctions, unique known (DA) junctions not found in GTEx, unique D, A, NDA junctions, and unique D, A, NDA junctions not found in GTEx per sample per cohort.These results were obtained using the all exonic (-E) and all intronic (-I) splice variant window parameters.

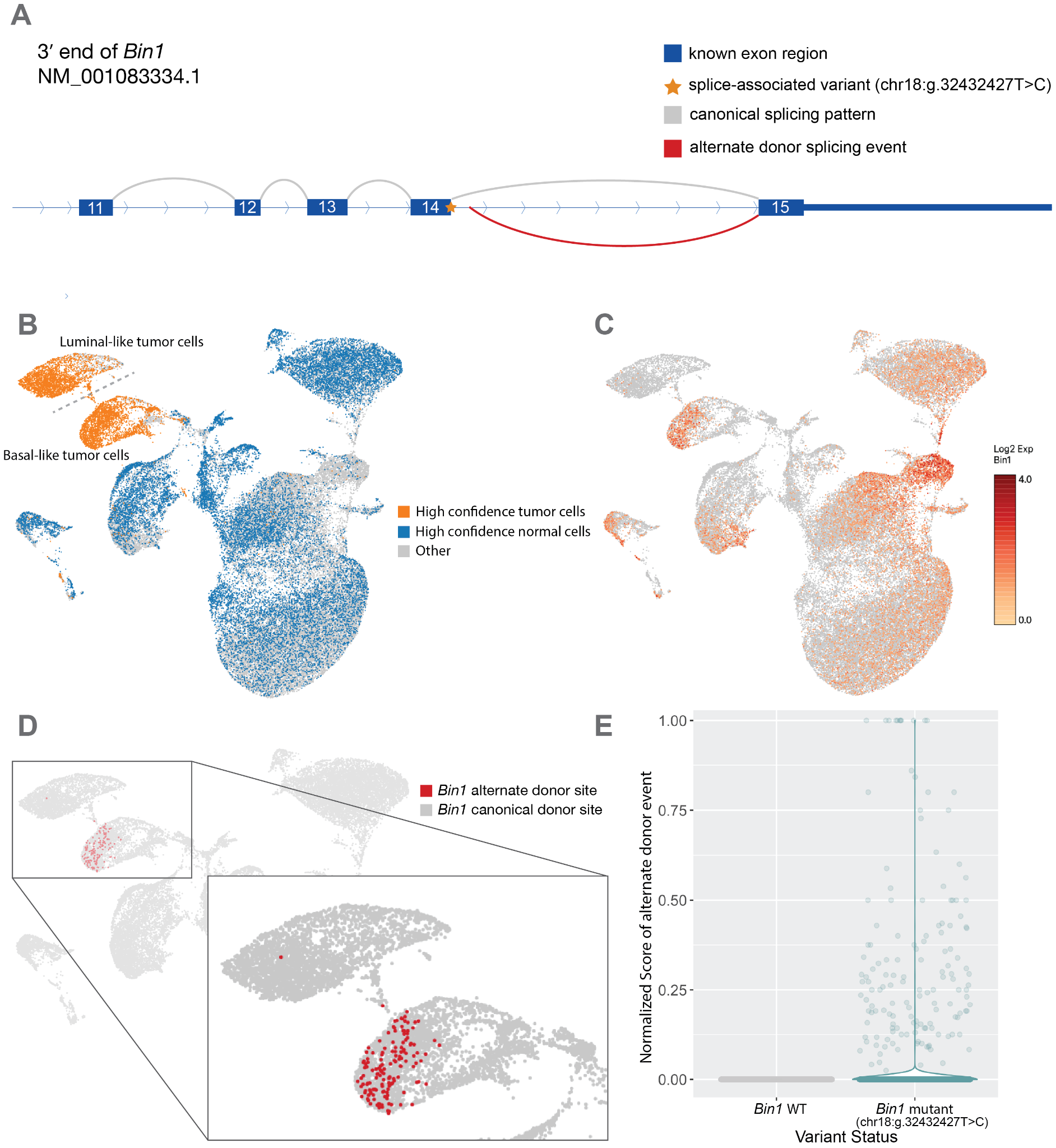

#### Supplementary Figure 5: Intronic SNV in *Bin1* associated with alternate donor usage by scRNA-seq analysis.

**A)** Schematic of a single nucleotide variant (mm10, chr18:g.32432427T>C, c.1516+2 position of intron 14 of transcript NM_001083334.1) within an intron of *Bin1*. This variant is significantly associated with an alternate donor event causing the formation of a novel junction. This result was found using the default splice variant window parameter (i2e3). **B)** UMAP projection of single cells from MCB6C organoid derived tumors with high confidence tumor cells (orange) and high confidence normal cells (blue) highlighted. **C)** UMAP projection of single cells from MCB6C organoid derived tumors overlaid with Log2 expression of Bin1. **D)** Zoomed view of cells containing the Bin1 alternate donor event. **E)** Violin plot comparing the normalized junction score of the novel alternate donor event in cells with and without the variant.

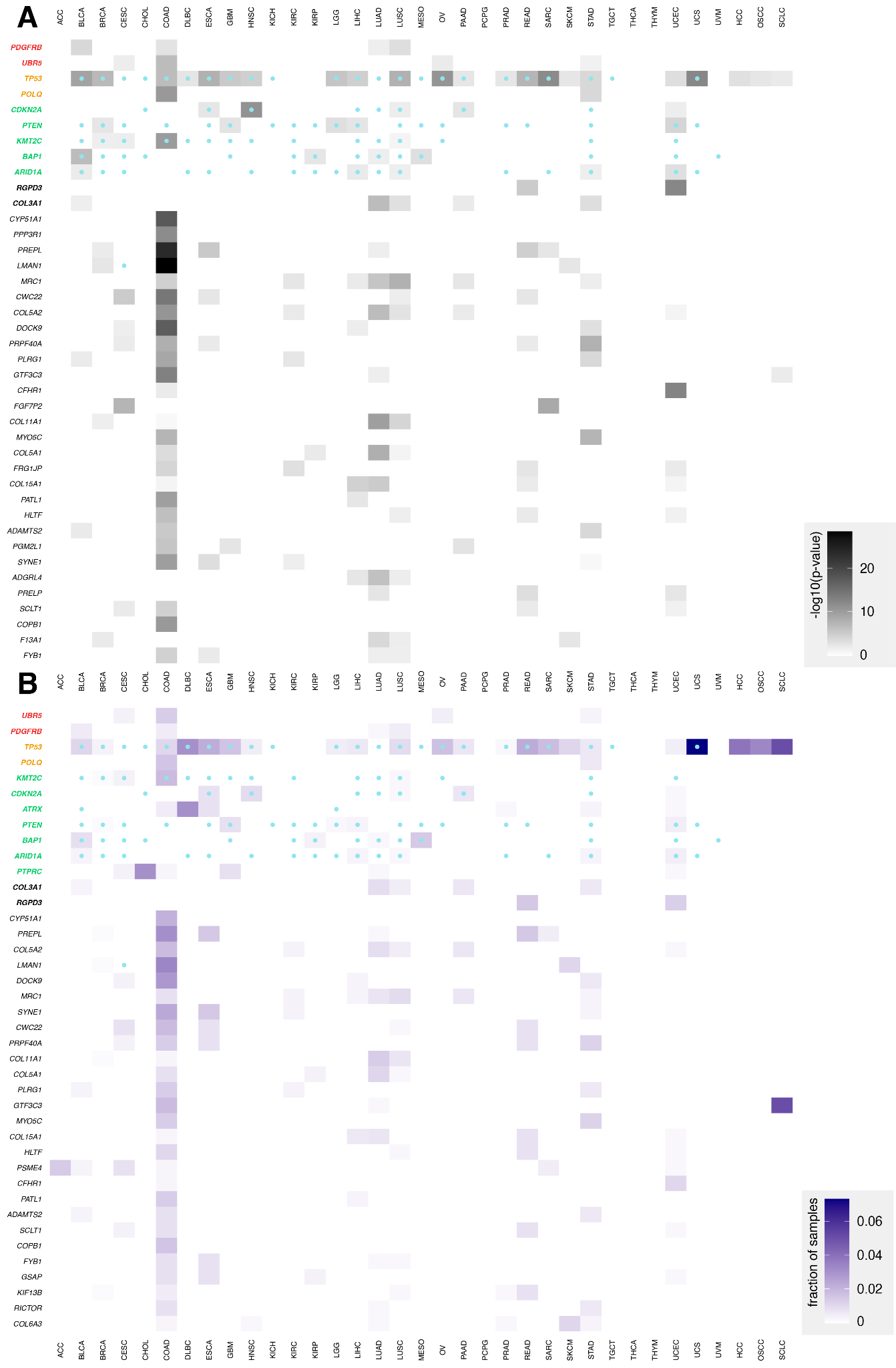

#### Supplementary Figure 6. Pan-cancer analysis of cohorts from TCGA and MGI reveals genes recurrently disrupted by variants which promote splicing of particular canonical junctions

Results of analysis for recurrently disrupted genes in each TCGA and MGI cohort. **A)** Rows correspond to the 40 most frequently recurring genes, as ranked by binomial p-value. Genes are clustered by whether they were annotated by the CGC as an oncogene (red), an oncogene and tumor suppressor gene (yellow), a tumor suppressor gene (green), or another type of cancer-relevant gene (black, bold). Shading corresponds to −log10(p value) and columns represent cancer types. Blue marks within cells indicate that the gene was annotated by CHASMplus as a driver within a given TCGA cohort. **B)** Rows correspond to the 40 most frequently recurring genes, as ranked by fraction of samples. Shading corresponds to the fraction of samples and columns represent cancer types. Blue dots within cells indicate that the gene was annotated by CHASMplus as a driver within a given TCGA cohort. These results were obtained using the default splice variant window parameter (i2e3).

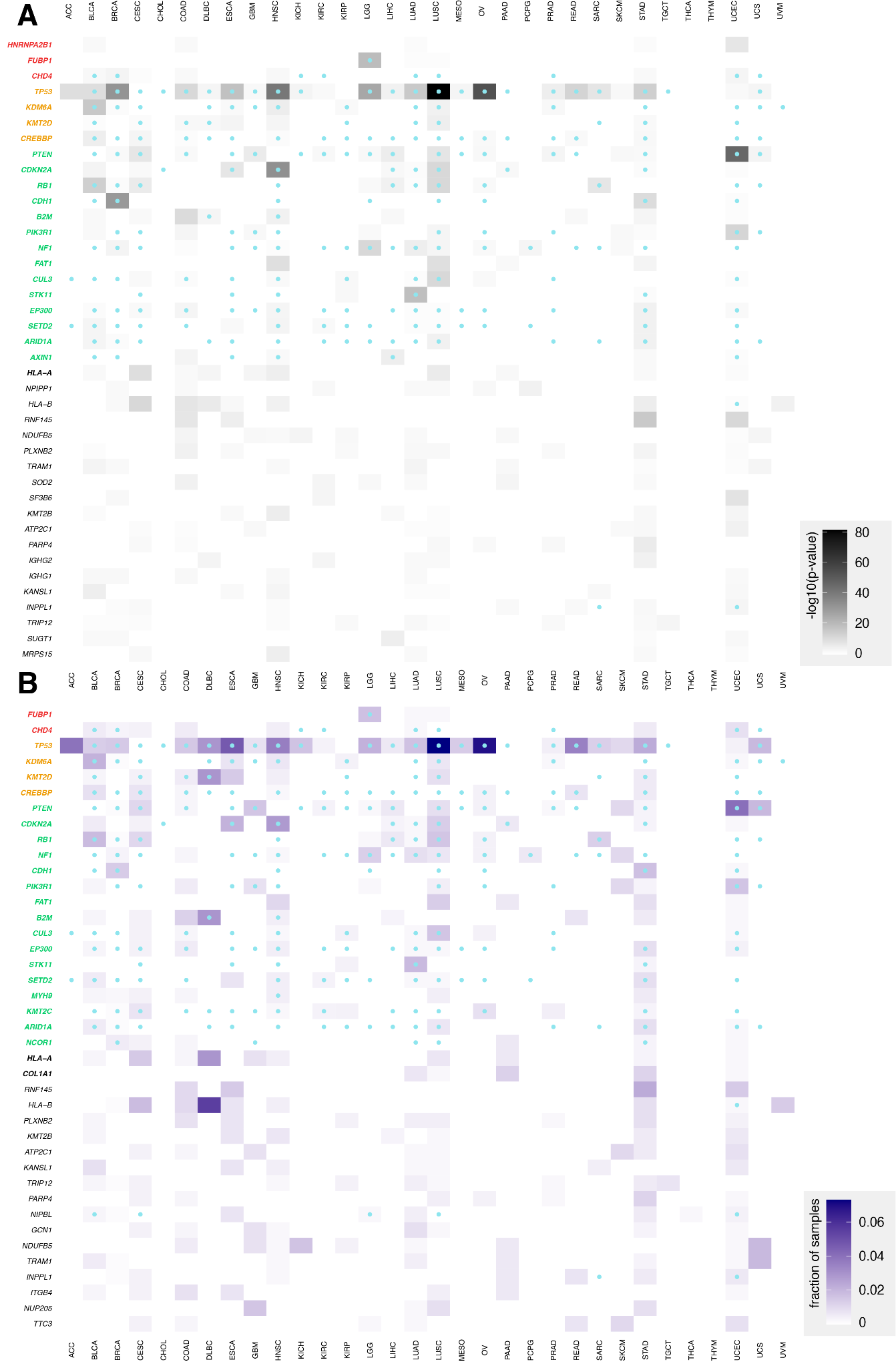

#### Supplementary Figure 7. TCGA pan-cancer analysis reveals genes recurrently disrupted by variants which cause non-canonical splicing patterns

Results of analysis for recurrently disrupted genes in each TCGA cohort. **A)** Rows correspond to the 40 most frequently recurring genes, as ranked by binomial p-value. Genes are clustered by whether they were annotated by the CGC as an oncogene (red), an oncogene and tumor suppressor gene (yellow), a tumor suppressor gene (green), or another type of cancer-relevant gene (black, bold). Shading corresponds to −log10(p value) and columns represent cancer types. Blue marks within cells indicate that the gene was annotated by CHASMplus as a driver within a given TCGA cohort. **B)** Rows correspond to the 40 most frequently recurring genes, as ranked by fraction of samples. Shading corresponds to the fraction of samples and columns represent cancer types. Blue dots within cells indicate that the gene was annotated by CHASMplus as a driver within a given TCGA cohort. These results were obtained using the default splice variant window parameter (i2e3).

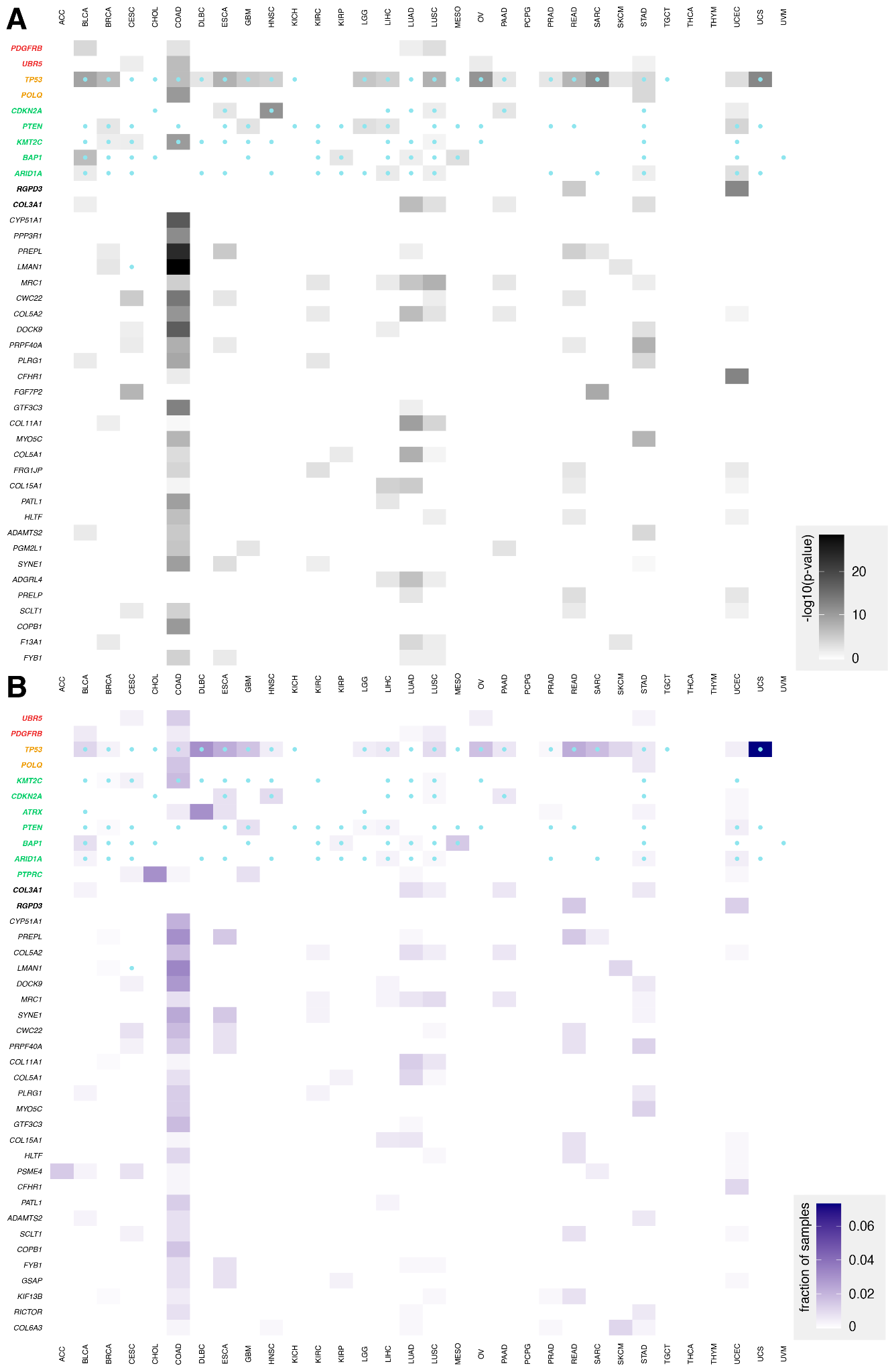

#### Supplementary Figure 8. TCGA pan-cancer analysis reveals genes recurrently disrupted by variants which promote splicing of particular canonical junctions

Results of analysis for recurrently disrupted genes in each TCGA cohort. **A)** Rows correspond to the 40 most frequently recurring genes, as ranked by binomial p-value. Genes are clustered by whether they were annotated by the CGC as an oncogene (red), an oncogene and tumor suppressor gene (yellow), a tumor suppressor gene (green), or another type of cancer-relevant gene (black, bold). Shading corresponds to −log10(p value) and columns represent cancer types. Blue marks within cells indicate that the gene was annotated by CHASMplus as a driver within a given TCGA cohort. **B)** Rows correspond to the 40 most frequently recurring genes, as ranked by fraction of samples. Shading corresponds to the fraction of samples and columns represent cancer types. Blue dots within cells indicate that the gene was annotated by CHASMplus as a driver within a given TCGA cohort. These results were obtained using the default splice variant window parameter (i2e3).

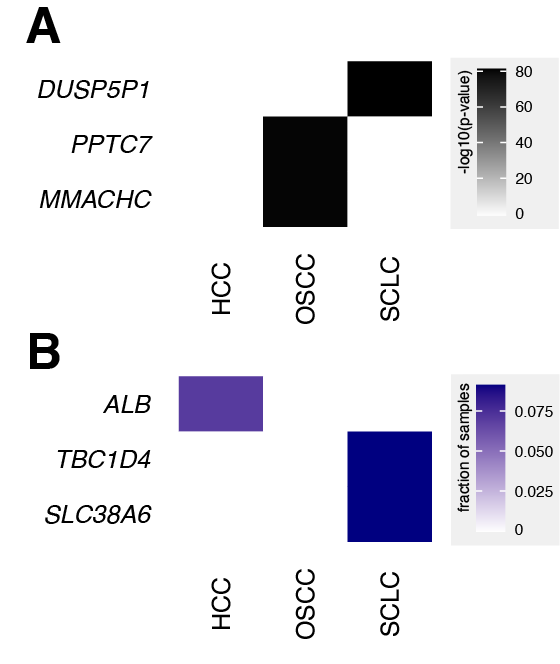

#### Supplementary Figure 9. Analysis of MGI cohorts reveals genes recurrently disrupted by variants which cause non-canonical splicing patterns

Results of analysis for recurrently disrupted genes in each MGI cohort. **A)** Rows correspond to the 40 most frequently recurring genes, as ranked by binomial p-value. Genes are clustered by whether they were annotated by the CGC as an oncogene (red), an oncogene and tumor suppressor gene (yellow), a tumor suppressor gene (green), or another type of cancer-relevant gene (black, bold). Shading corresponds to −log10(p value) and columns represent cancer types. **B)** Rows correspond to the 40 most frequently recurring genes, as ranked by the fraction of samples. Shading corresponds to the fraction of samples and columns represent cancer types. These results were obtained using the default splice variant window parameter (i2e3).

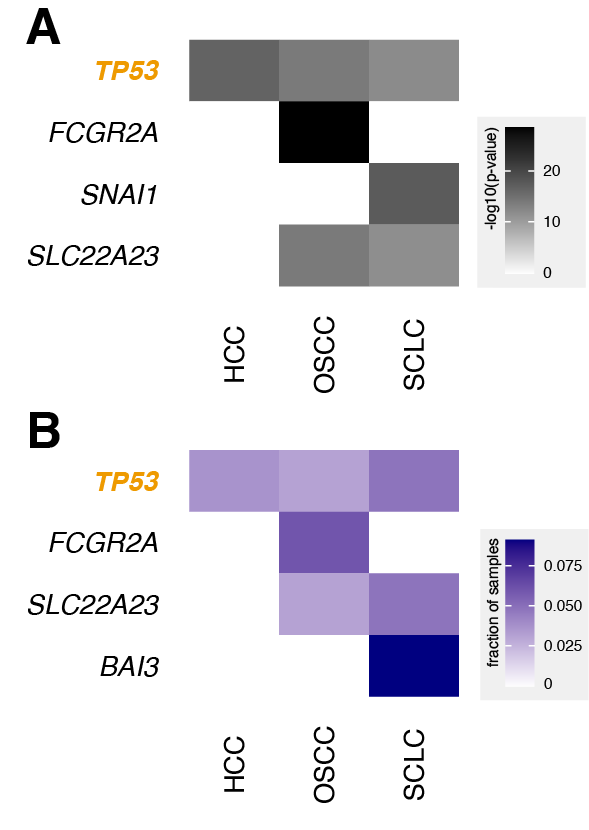

#### Supplementary Figure 10. Analysis of MGI cohorts reveals genes recurrently disrupted by variants which promote splicing of particular canonical junctions

Results of analysis for recurrently disrupted genes in each MGI cohort. **A)** Rows correspond to the 4 most frequently recurring genes, as ranked by binomial p-value. Genes are clustered by whether they were annotated by the CGC as an oncogene (red), an oncogene and tumor suppressor gene (yellow), a tumor suppressor gene (green), or another type of cancer-relevant gene (black, bold). Shading corresponds to −log10(p value) and columns represent cancer types. **B)** Rows correspond to the 4 most frequently recurring genes, as ranked by the fraction of samples. Shading corresponds to the fraction of samples and columns represent cancer types. These results were obtained using the default splice variant window parameter (i2e3).

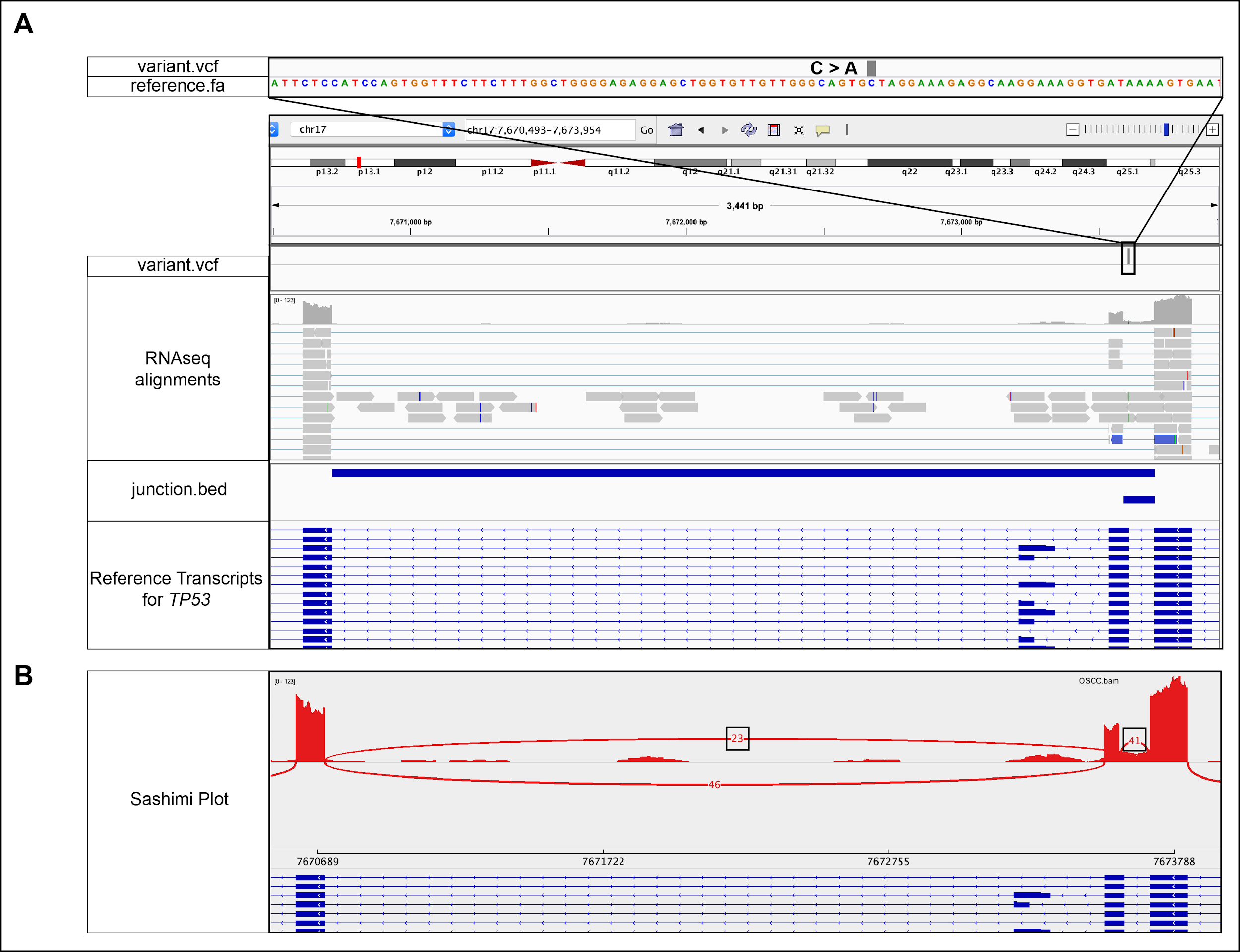

#### Supplementary Figure 11: Intronic SNV in *TP53* associated with exon skipping and alternative donor usage.

**A)** IGV snapshot of a single nucleotide variant (GRCh38, chr17:g.7673609C>A, NM_000546.5) within an intron of *TP53* in an OSCC sample. This variant is associated with skipping of exon 9 (23 reads of support) and use of an alternate acceptor site (41 reads of support). This result was found using the default splice variant window parameter (i2e3). **B)** Sashimi plot visualization of the novel junction.

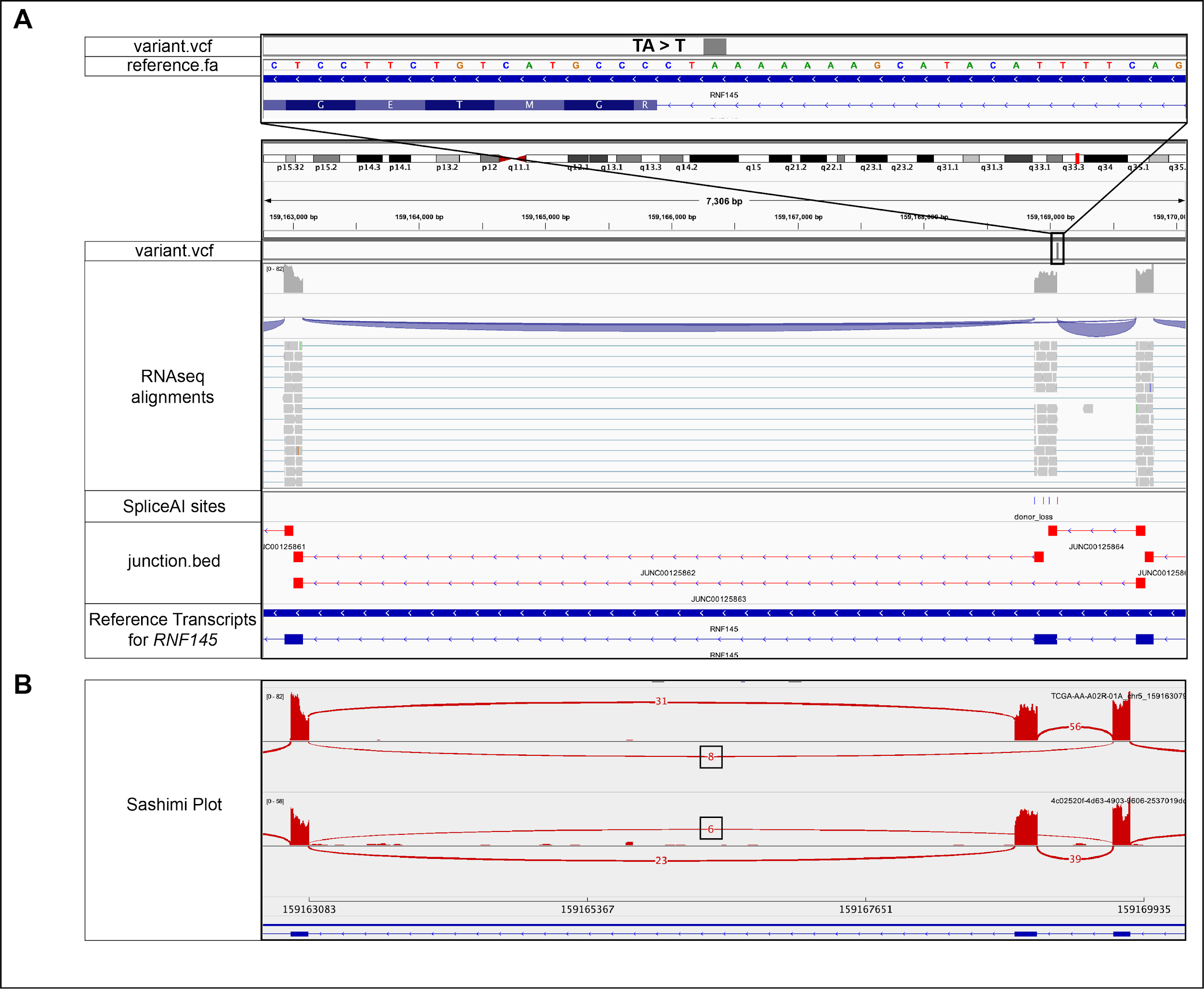

#### Supplementary Figure 12: Intronic deletion in *RNF145* associated with skipping of exon 8.

**A)** IGV snapshot of a single nucleotide variant (GRCh38, chr5:g.159169058delA, NM_001199383.2) within an intron of *RNF145* in COAD samples. This variant is associated with an exon skipping event with 8 and 6 reads of support for the samples shown. This result was found using the default splice variant window parameter (i2e3). **B)** Sashimi plot visualization of the novel junction.

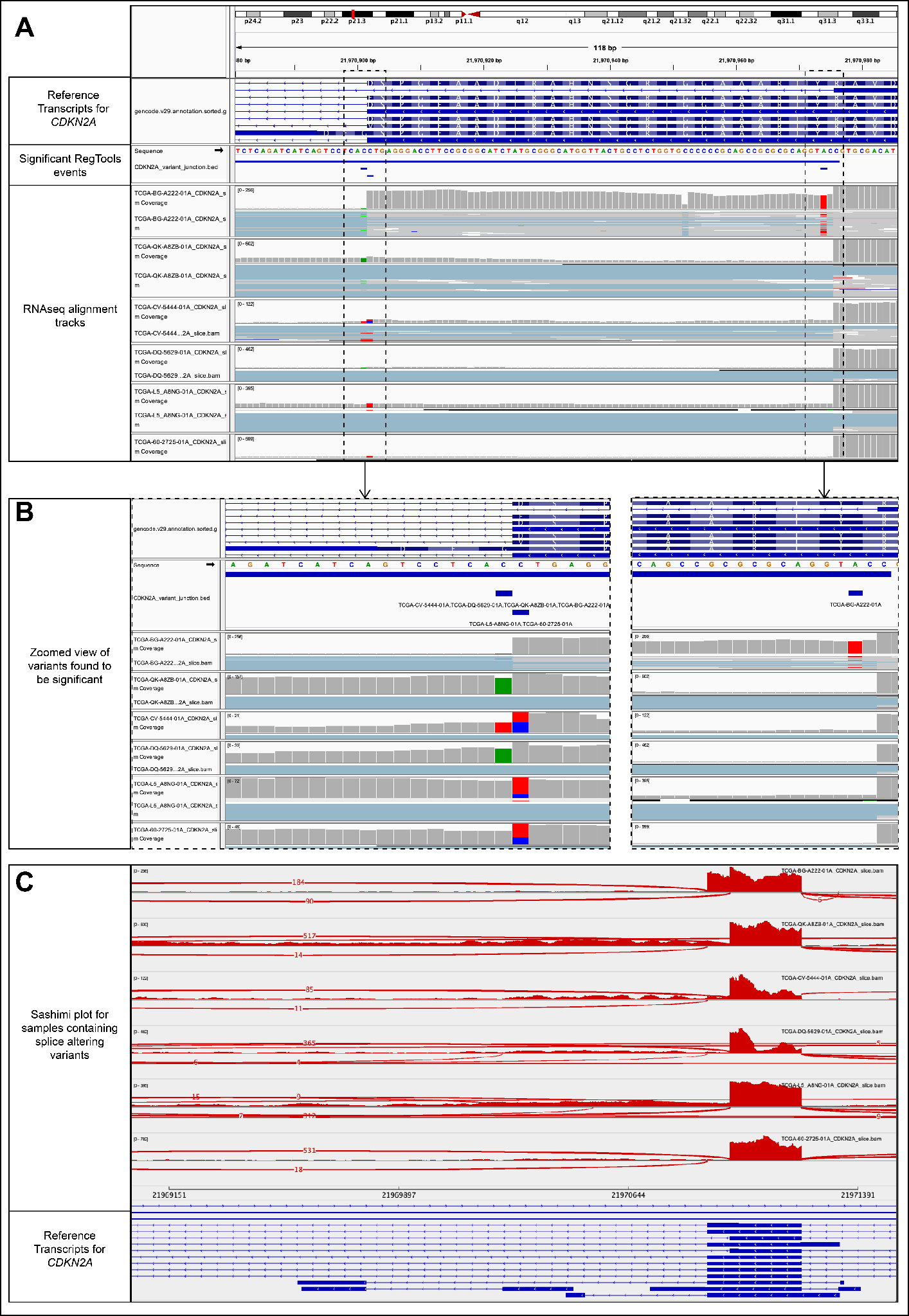

#### Supplementary Figure 13: Several SNVs in *CDKN2A* associated with alternate donor usage leading to an alternate known junction that may cause C-terminus modification.

**A)** IGV snapshot of three variant positions in *CDKN2A* found to be associated with usage of an alternate donor site within exon 2 that leads to the formation of an alternate known junction included in an alternate transcript, ENST00000579122.1. This result was found using the default splice variant window parameter (i2e3) for known (DA) junctions. **B)** Zoomed in view of the variants identified by RegTools that are associated with alternate donor usage. Two of these variant positions flank the donor site that is no longer being used. **C)** Sashimi plot visualizations for samples containing the identified variants that show alternate donor usage.

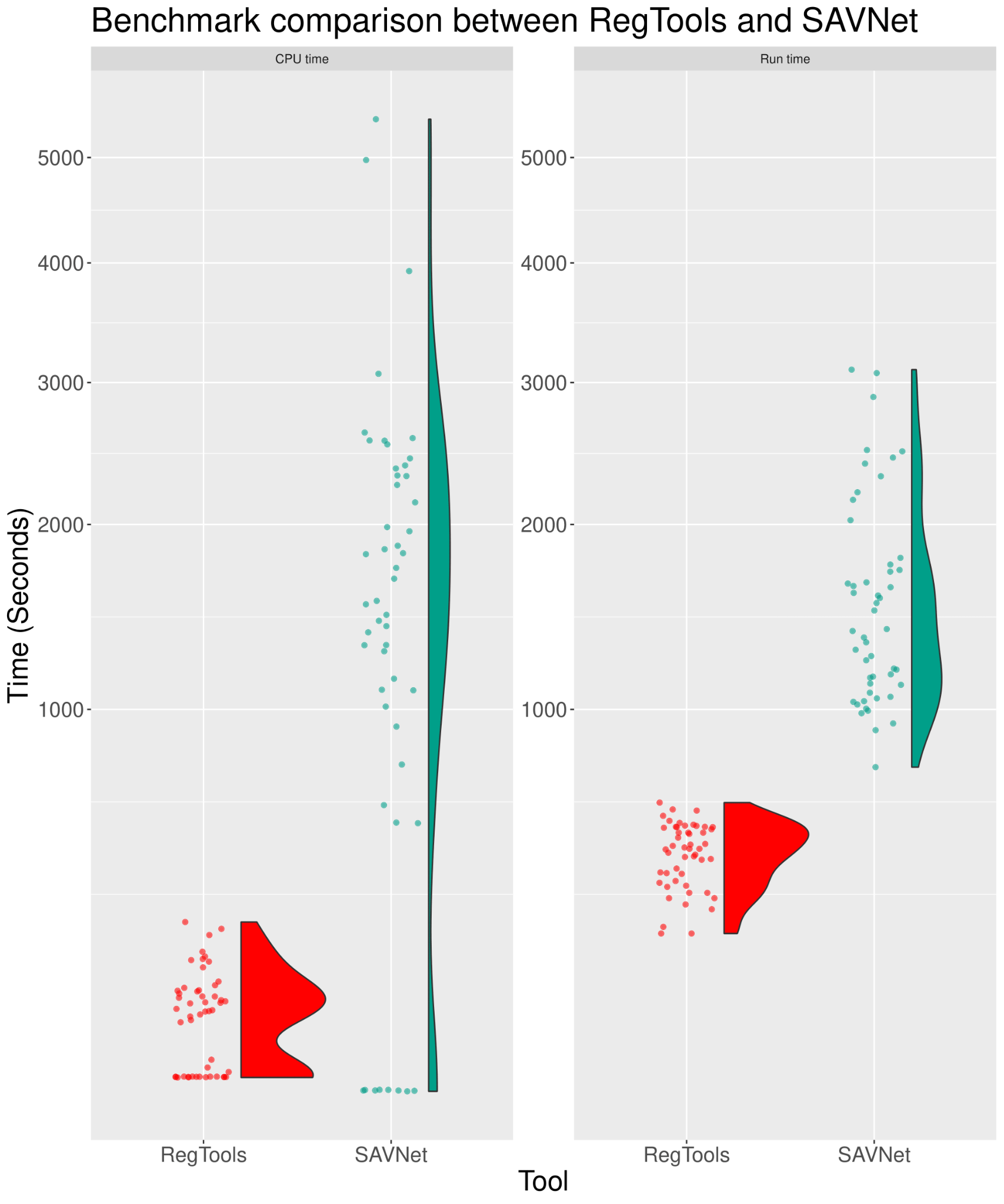

#### Supplementary Figure 14. Benchmarking of RegTools and SAVNet

The total CPU time (System Time + User Time) and real-time are plotted against both RegTools and SAVNet for fifty LUAD samples from TCGA. Each sample is represented as a point and a half violin plot is plotted alongside the dot plot to show the distribution of runtimes. These results were obtained using the default splice variant window parameter (i2e3).
